# Small-world scale-free brain networks from EEG with application to motor imagery decoding and brain fingerprinting

**DOI:** 10.1101/2025.04.17.649421

**Authors:** Mohammad Davood Khalili, Vahid Abootalebi, Hamid Saeedi-Sourck, Andrea Santoro, Harry H. Behjat

**Affiliations:** Department of Electrical Engineering, Yazd University, Yazd, Iran; Neuro-X Institute, École Polytechnique Fédérale de Lausanne, Geneva, Switzerland; ISI Foundation, Turin, Italy; Department of Biomedical Engineering, Lund University, Lund, Sweden; Department of Clinical Sciences Malmö, Lund University, Lund, Sweden

**Keywords:** EEG, small-world, scale-free, graph reduction, graph signal processing, motor imagery, brain fingerprinting

## Abstract

Developing individualized spatial models that capture the complex dynamics of multi-electrode EEG data is essential for accurately decoding global neural activity. A widely used approach is network modeling, where electrodes are represented as nodes. A key challenge lies in defining the network edges and weights, as precise connectivity estimation is critical for enhancing neural characterization and extracting discriminative features, such as those needed for task decoding. Traditional EEG-derived brain graphs often fail to capture biologically grounded organizational principles such as small-world structure and heavy-tailed (scale-free) connectivity patterns. To address this gap, we introduce a framework for inferring subject-specific EEG-based brain graphs that explicitly designed to exhibit small-world and scale-free properties. Our approach begins by computing phase-locking values (PLV) between EEG channel pairs to build a backbone graph, which is then refined into an individualized small-world and scale-free network. To reduce computational complexity while preserving subject-specific characteristics, we apply Kron reduction to the resulting graph. Using two public EEG datasets, we evaluate the proposed method on motor imagery (MI) decoding and brain fingerprinting tasks. Our approach improves MI classification accuracy by 4–7% compared to conventional PLV, small-world, and scale-free graph models, and enhances differential identifiability in fingerprinting by 8–20% across six canonical frequency bands. These gains were statistically significant in both applications. Moreover, integrating graph signal processing features derived from our constructed graphs with classical EEG features further boosts performance. Overall, our findings highlight the potential of the proposed graph construction framework to enhance EEG analysis. By jointly capturing local segregation, global integration, and hub-driven hierarchical organization, the method strengthens downstream decoding and identification tasks, with promising implications for a wide range of applications in cognitive neuroscience and brain-computer interface research.

## 1. Introduction

Electroencephalography (EEG), which records neural activity via a network of scalp electrodes, remains a cornerstone technique for probing brain dynamics due to its millisecond temporal resolution and non-invasiveness [1]. However, its relatively low spatial resolution and the presence of volume conduction frequently introduce high inter-channel correlations, complicating the interpretation of functional interactions [2]. In recent years, network science has emerged as a powerful paradigm for elucidating the organizational principles of brain architecture and its complex functional interactions [3-5]. Within this framework, brain regions or sensors are represented as nodes, while their structural or statistical relationships are encoded as weighted edges [6, 7]. Nevertheless, constructing a biologically meaningful, and computationally tractable graph representation of EEG data remains an open and challenging research problem [8-10], critical for advancing our understanding of brain connectivity and improving downstream analytical tasks.

Empirical evidence shows that the brain has a small-world modular structure [4], comprising canonical functional networks across diverse temporal and spatial scales [11], alongside structural and functional hubs. This has significant biological implications, equipping the brain with an infrastructure to maintain the equilibrium between integration and segregation [4]. Recent studies suggest that this optimal trade-off stems from an evolutionary process aimed at minimizing wiring costs while maximizing global information processing efficiency [12-14]. Theoretically, it is well-accepted that the brain optimizes its operations by minimizing the costs of information processing, maximizing efficiency, and adaptively rerouting its operations to ensure resilience [15-17].

While EEG-based functional graphs have been widely employed—using, namely, correlation, coherence, PLV, partial correlation, or data-driven connectivity estimation—most existing approaches suffer from several limitations. Thresholded connectivity graphs often fail to reproduce small-world structure; graph learning methods frequently ignore scale-free behavior; and unconstrained data-driven techniques may generate networks that are inconsistent across subjects, sensitive to noise, or limited in neurobiological interpretability. Moreover, prior work typically focuses solely on estimating connectivity strengths, without explicitly enforcing topological properties known to characterize real neural systems. This leads to EEG-derived graphs that may be analytically useful but remain biologically ungrounded.

These diverse requirements underscore the need for EEG graphs that balance biological plausibility with computational tractability. In response, we propose a novel class of EEG-based brain graphs, referred to as Small-world Scale-free (SWSF), along with a lightweight variant incorporating Kron reduction, denoted SWSF_Kron_. The proposed Small-world Scale-free brain graphs integrate small-world and scale-free characteristics, offering an effective balance between high communication efficiency and minimal wiring cost. In the SWSF graphs, a small-world architecture encodes global connectivity among a set of high-significance nodes, identified through functional clustering, while a scale-free structure models local connectivity among lower-significance nodes surrounding these hubs. The variant denoted SWSF_Kron_, incorporates Kron reduction as a post-processing step, reducing computational complexity while preserving (1) inter-regional functional connectivity patterns, (2) spectral characteristics, and (3) essential topological features, including small-worldness and scale-freeness, through the Schur complement. Brain graphs exhibiting these features are known for their efficiency and resilience, which are critical in shaping plasticity, cognition, and insights into neurological diseases and disorders. Both the SWSF and SWSF_Kron_ models accurately capture the spatial organization of the brain, emphasizing short transmission paths for high efficiency, a high clustering coefficient, and the emergence of hubs that facilitate coordinated information flow. In summary, these models effectively reflect the brain’s balance between integration and segregation, establishing themselves as principled and biologically grounded EEG-based brain graphs. To demonstrate the applicability of the proposed EEG-based brain graphs, we explore two proof-of-concept applications: motor imagery (MI) decoding [18-20] and brain fingerprinting [21-23]. Accurately distinguishing mental states from EEG data remains a persistent challenge, with a continued wide array of methods being proposed to address this issue [24]. Some methods focus on time- and frequency-based feature [25-27], while others focus on deriving spatial features from EEG topographies [28, 29]. Recent advances in deep learning and clinical EEG analysis also underscore the need for principled graph representations, as demonstrated in work on multiscale spatial–temporal MI decoding [30] and BCI-based prognosis in cognitive motor dissociation [31]. Using two open-access EEG datasets, we show that our SWSF and SWSF_Kron_ approaches provide superior performance over small-world and scale-free brain graphs in MI decoding and brain fingerprinting.

The remainder of the paper is organized as follows: Section 2 overviews key GSP concepts, details the proposed SWSF brain graph design, describes the datasets, and outlines methods for MI decoding and brain fingerprinting. Section 3 presents and discusses the experimental results. Section 4 provides our concluding remarks.

## 2. Materials and Methods

Table 1 summarizes the notation used in this paper.

**Table 1:**
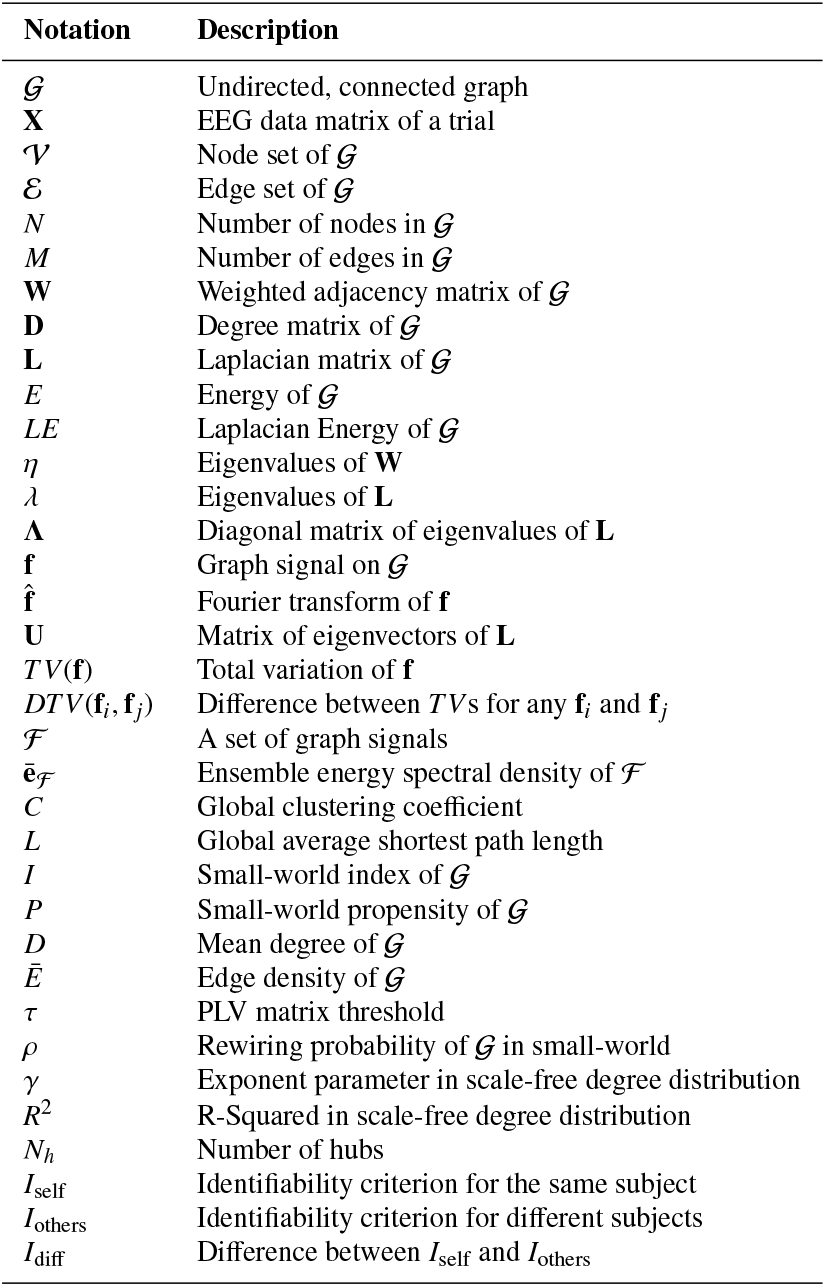
List of the main notations.

| Notation | Description |
| --- | --- |
| $\mathcal{G}$ | Undirected, connected graph |
| $\mathbf{X}$ | EEG data matrix of a trial |
| $\mathcal{V}$ | Node set of $\mathcal{G}$ |
| $\mathcal{E}$ | Edge set of $\mathcal{G}$ |
| $N$ | Number of nodes in $\mathcal{G}$ |
| $M$ | Number of edges in $\mathcal{G}$ |
| $\mathbf{W}$ | Weighted adjacency matrix of $\mathcal{G}$ |
| $\mathbf{D}$ | Degree matrix of $\mathcal{G}$ |
| $\mathbf{L}$ | Laplacian matrix of $\mathcal{G}$ |
| $E$ | Energy of $\mathcal{G}$ |
| $LE$ | Laplacian Energy of $\mathcal{G}$ |
| $\eta$ | Eigenvalues of $\mathbf{W}$ |
| $\lambda$ | Eigenvalues of $\mathbf{L}$ |
| $\Lambda$ | Diagonal matrix of eigenvalues of $\mathbf{L}$ |
| $\mathbf{f}$ | Graph signal on $\mathcal{G}$ |
| $\hat{\mathbf{f}}$ | Fourier transform of $\mathbf{f}$ |
| $\mathbf{U}$ | Matrix of eigenvectors of $\mathbf{L}$ |
| $TV(\mathbf{f})$ | Total variation of $\mathbf{f}$ |
| $DTV(\mathbf{f}_i, \mathbf{f}_j)$ | Difference between $TV$s for any $\mathbf{f}_i$ and $\mathbf{f}_j$ |
| $\mathcal{F}$ | A set of graph signals |
| $\bar{e}_{\mathcal{F}}$ | Ensemble energy spectral density of $\mathcal{F}$ |
| $C$ | Global clustering coefficient |
| $L$ | Global average shortest path length |
| $I$ | Small-world index of $\mathcal{G}$ |
| $P$ | Small-world propensity of $\mathcal{G}$ |
| $D$ | Mean degree of $\mathcal{G}$ |
| $\bar{E}$ | Edge density of $\mathcal{G}$ |
| $\tau$ | PLV matrix threshold |
| $\rho$ | Rewiring probability of $\mathcal{G}$ in small-world |
| $\gamma$ | Exponent parameter in scale-free degree distribution |
| $R^2$ | R-Squared in scale-free degree distribution |
| $N_h$ | Number of hubs |
| $I_{\text{self}}$ | Identifiability criterion for the same subject |
| $I_{\text{others}}$ | Identifiability criterion for different subjects |
| $I_{\text{diff}}$ | Difference between $I_{\text{self}}$ and $I_{\text{others}}$ |

### 2.1 Graphs, their spectra, and graph signals

Let 𝒢 = (𝒱, ℰ, **W**) denote an undirected connected graph characterized by the set of nodes, 𝒱 where |𝒱| = *N* is the number of nodes and the set of edges ℰ, where |ℰ| = *M* is the number of edges, and an *N* × *N* weighted adjacency matrix **W**, with elements *W*_*i, j*_ >= 0 denoting the weight of edge (*i, j*) ∈ ℰ. The combinatorial graph Laplacian matrix is defined as **L** = **D** − **W**, where **D** is the diagonal degree matrix with elements 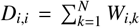. The energy of 𝒢 is defined as 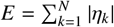, where {η_*k*_} denote the eigenvalues of **W**.

The Laplacian energy of G is defined as 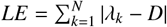 [34], where *D* is the average node degree, {λ_*k*_} denote the eigenvalues of **L** obtained as diagonal elements of Λ in **L** = **U**Λ**U**^*T*^, with **U** denoting the corresponding eigenvectors.

Let **f** ∈ ℝ^*N*^ denote a graph signal on 𝒢, whose *n*-th component represents the signal value at the *n*-th node of 𝒢. The graph Fourier transform (GFT) of **f** is given as [35]:

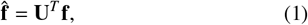

which satisfies the Parseval’s relation: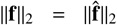, ensuring energy conservation between the node and spectral domain.

The total variation of **f** is defined as:

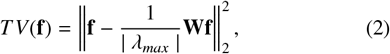

where ‖ · ‖_2_ denotes the Frobenius norm and λ_*max*_ is the largest-magnitude eigenvalue of **W** [37]. Using *TV*(**f**), the difference between the *TV* of the *i*-th and *j*-th graph signal is given as [38, 39]:

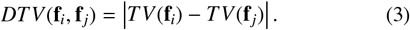

Lastly, the ensemble energy spectral density of a given graph signal set 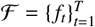 is obtained as [36]:

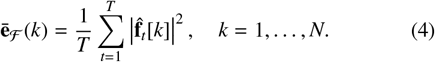

### 2.2. Network Models

Multiple network models have been proposed to represent the structural properties found in real-world networks [40]. Erdõs-Rényi random graphs are networks wherein each edge has a fixed probability of being present or absent, resulting in a Poisson degree distribution [41]. They have a small clustering coefficient (CC) and characteristic path length (CPL) [42-44], and thus are not suitable to represent most real-world networks characterized by the presence of modules, hubs, and high CC. In contrast, regular networks feature structured link patterns—such as rings or grids—where each node connects to the same number of neighbors. These lattices typically have higher clustering coefficients and longer characteristic path lengths than random graphs [4].

Barabási and Albert [45] introduced a network growth model with preferential attachment, where new nodes connect to existing ones based on their degree, favoring high-degree nodes. Over many iterations, this forms a scale-free (SF) network with a power-law degree distribution (∼3) and prominent hubs [46]. In contrast, Watts and Strogatz [47] proposed the seminal small-world (SW) network model, which begins with a regular ring lattice where each node connects to its *D*/2 nearest neighbors clockwise and *D*/2 counterclockwise. This initial configuration exhibits three defining properties: high CC, long CPL, and absence of hubs. The core mechanism of the model is stochastic edge rewiring: each existing edge is randomly reconfigured with probability ρ, producing three topological regimes:

- ρ = 0: Purely regular lattice (high CC, long CPL)
- 0 < ρ < 1: SW network (high CC, short CPL)
- ρ = 1: Fully randomized network (low CC, short CPL).

For EEG applications, we generalize this model to 2D scalp-projected electrode arrays. Connections are defined relative to each electrode’s radial line to the geometric centroid, with *D*/2 nearest neighbors selected in each of the clockwise and counterclockwise angular half-planes. This adaptation preserves the SW generation mechanism through angular rewiring, respects neuroanatomical constraints, and maintains functional balance: high CC for local specialization and short CPL for global integration [13].

Let *C* and *L* denote the CC and CPL of 𝒢, respectively. Let *C*^(rnd)^ and *C*^(reg)^ denote the CC of a random and regular graph of size 𝒢, respectively. Similarly, let *L*^(rnd)^ and *L*^(reg)^ denote the CPL of a random and regular graph of size 𝒢, respectively. The extent of small-worldness of 𝒢 can be quantified using the SW index (*I*) measure, defined as [48]:

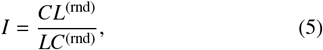

or the SW propensity (*P*) measure, defined as [49]:

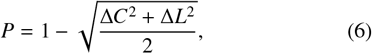

where Δ*C* and Δ*L* are given as:

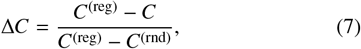

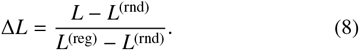

In a SW network [47], *C*/*C*^(rnd)^ > 1 and *L*/*L*^(rnd)^ ≈ 1, thus, a scalar summary of a small-world graph is *I* > 1 or *P* > 0.6 [48, 49]. The sparsity of the graph, which is a key parameter in its construction, can be quantified using the edge density, denoted *Ē*, and computed as *Ē* = 2*M*/*N*(*N* − 1) [17].

### 2.3. Proposed Method: SWSF Brain Graph Design

An overview of the proposed pipeline for designing SWSF brain graphs from EEG is illustrated as a block diagram in Fig. 1. The pipeline takes as input an *N*-channel EEG recording, consisting of *K* trials, each of length *T*, represented as a matrix *N* × *T*, denoted **X**. The time courses are assumed to have been suitably bandpass filtered to be confined in a desired frequency range. First, we construct a phase locking value (PLV) graph [50] that quantifies the phase synchronization of the electrodes, a widely used measure in brain-computer interface (BCI) studies [51-53]. Functional connectivity was quantified using PLV, which estimates the consistency of phase differences between pairs of neural signals. PLV provides a frequency-specific and temporally precise measure of oscillatory synchrony, capturing functional coupling rather than amplitude co-modulation.

**Fig. 1:**
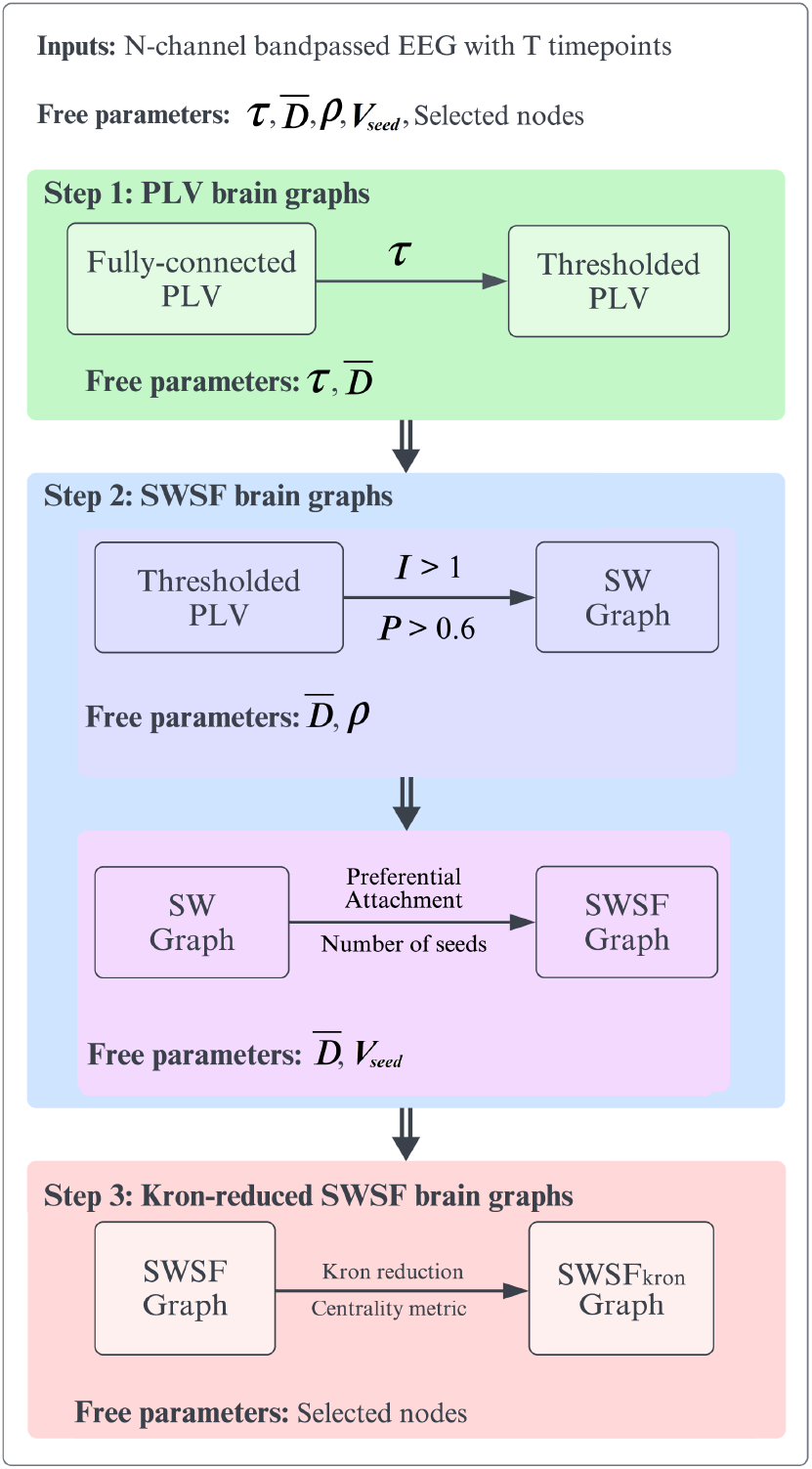
The block diagram of the proposed method for inferring an SWSF graph from a given set of EEG data.

The analytic signal for each row *n* (denoted **X**_*n*,:_) is computed using the Hilbert Transform, denoted **X**^(*a*)^, and is composed of real (**X**^(*r*)^ = **X**) and imaginary (**X**^(*i*)^) components, i.e., **X**^(*a*)^ = **X**^(*r*)^ + *i***X**^(*i*)^ [54].^1^ An instantaneous phase matrix, denoted **X**^(*θ*)^, is then constructed, with elements:

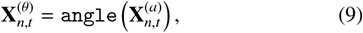

representing the phase of electrode *n* at time point *t*. Using **X**^(*θ*)^, we computed a PLV graph per trial, defined by its weighted adjacency matrix, a symmetric *N × N* matrix, denoted **W**, with elements **W**_*n,m*_ ∈ [0, 1] defined as the ensemble-averaged instantaneous PLV between pairs of electrodes *n* and *m*:

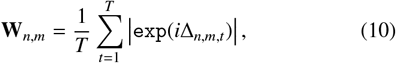

where 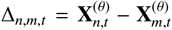. If electrodes *n* and *m* have similar phases across time, **W**_*n,m*_ →1 whereas if they have orthogonal phases **W**_*n,m*_ → 0. PLV matrices were interpreted as functional networks from which we derived small-world metrics, including the clustering coefficient, characteristic path length, and small-worldness index. These metrics quantify the balance between local segregation and global integration, two complementary and well-established principles of efficient brain organization. Inspired by prior works [52, 55, 56], the PLV graph was then sparsified by applying a threshold, denoted *τ* ∈ (0, 1), resulting in a thresholded PLV graph, denoted **W** ^(*τ*)^. We treat *τ* as a free parameter.

The methodologies for constructing small-world (SW), scale-free (SF), and small-world scale-free (SWSF) graph models are detailed in Algorithms 1, 2, and 3, respectively. These algorithms outline the systematic procedures for generating graph structures tailored to capture the topological characteristics critical to modeling brain networks.

Scale-free (SF) and small-world (SW) properties were treated as independent descriptors of network topology. SF characterizes the degree distribution (presence of hubs), whereas SW reflects the balance between local clustering and short path lengths. While hubs in a scale-free network can enhance small-world efficiency, small-world organization itself does not directly alter the degree distribution underlying heavy-tailed SF behavior.

The construction of SWSF graphs follows a two-step procedure. First, SW graphs are constructed by connecting each electrode to its spatially nearest neighbors in the 2D scalp-layout electrode array. The seed vertex set *V*_seed_ is then determined using application-specific functional clustering, selecting nodes with the highest degree centrality within each cluster. Subsequently, SF graphs are constructed using *V*_seed_ as the initial seed vertices. The SWSF graphs are derived from the thresholded weight matrix **W**^(*τ*)^, by selecting a suitable *D* as the construction parameter to balance computational complexity and network connectivity. To address the stochastic nature of the rewiring process, *N*_*b*_ bootstrap iterations were performed across all analyses: *N*_*b*_ = 50 for motor imagery decoding, and *N*_*b*_ = 30 for brain fingerprinting.

Random networks serve as a baseline for comparing the properties of SW networks. Thus, we used random networks during the construction of SW and SWSF graphs. Specifically, we constructed random networks with the desired average degree (*D*) and degree distributions identical to those of the original networks [57]. The parameters *C*^(rnd)^ and *L*^(rnd)^ were calculated as the means over *N*_*b*_ random graphs, following established methodologies in neuroscience [48, 58, 59]. Given 118 nodes in MI decoding, 1000 rewiring steps were sufficient to generate random graphs while preserving the nodal degree. This process was essential to establish the null distribution.

#### Algorithm 1

Small-World Graph Design

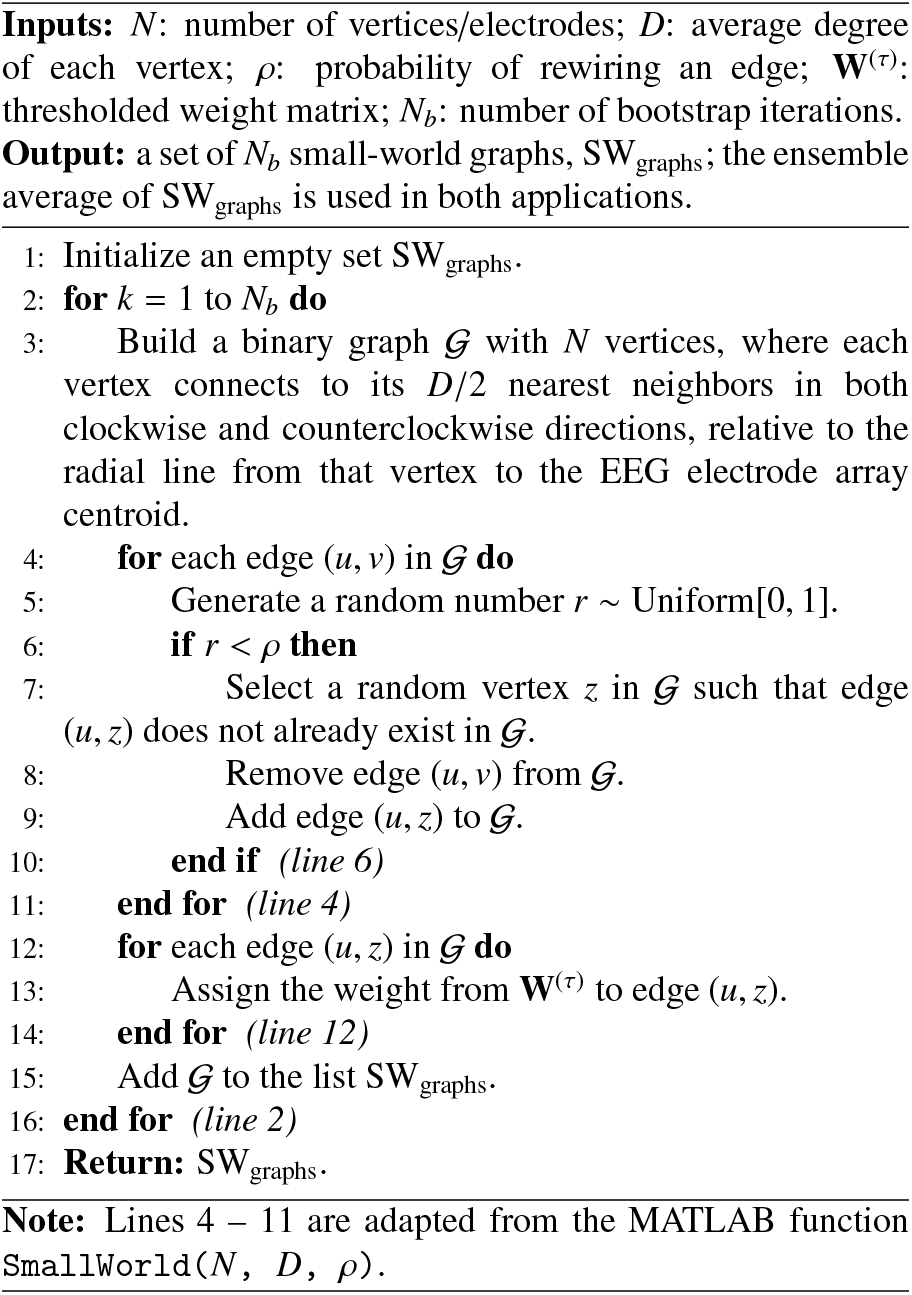

In step 10 of Algorithm 2 (Scale-free Graph Design), the probability of selecting an existing vertex *v*_*i*_ is given by *P*(*v*_*i*_) *=* deg(*v*_*i*_)*/* _*v*∈𝒢_ deg(*v*). This implements the preferential attachment principle, where nodes with larger degree have a proportionally higher likelihood of receiving new edges. This mechanism drives the emergence of hub nodes and produces a power-law degree distribution characteristic of scale-free networks. The probability of selecting any given node is therefore determined solely by its relative degree within the current graph.

To enhance robustness during the construction of SW, SF, and SWSF graphs while accounting for their stochastic nature, *N*_*b*_ brain graphs were generated for each scheme and each subject. Furthermore, we applied Kron reduction [60, 61] to reduce the size of the SWSF graphs. This reduction involved two steps: first, selecting the top 20% (i.e., 24 nodes) based on weighted nodal degrees, and second, simplifying the graph using Kron reduction with these selected nodes. Cross-validation results validating this node selection process (24 nodes) to construct the Kron-reduced SWSF graphs are shown in Fig. S1 (bottom, right) in the Supplementary Material. For both datasets, the Kron-reduced graph was constructed using a set of 24 retained nodes to maintain consistency and comparability across analyses.

#### Algorithm 2

Scale-Free Graph Design

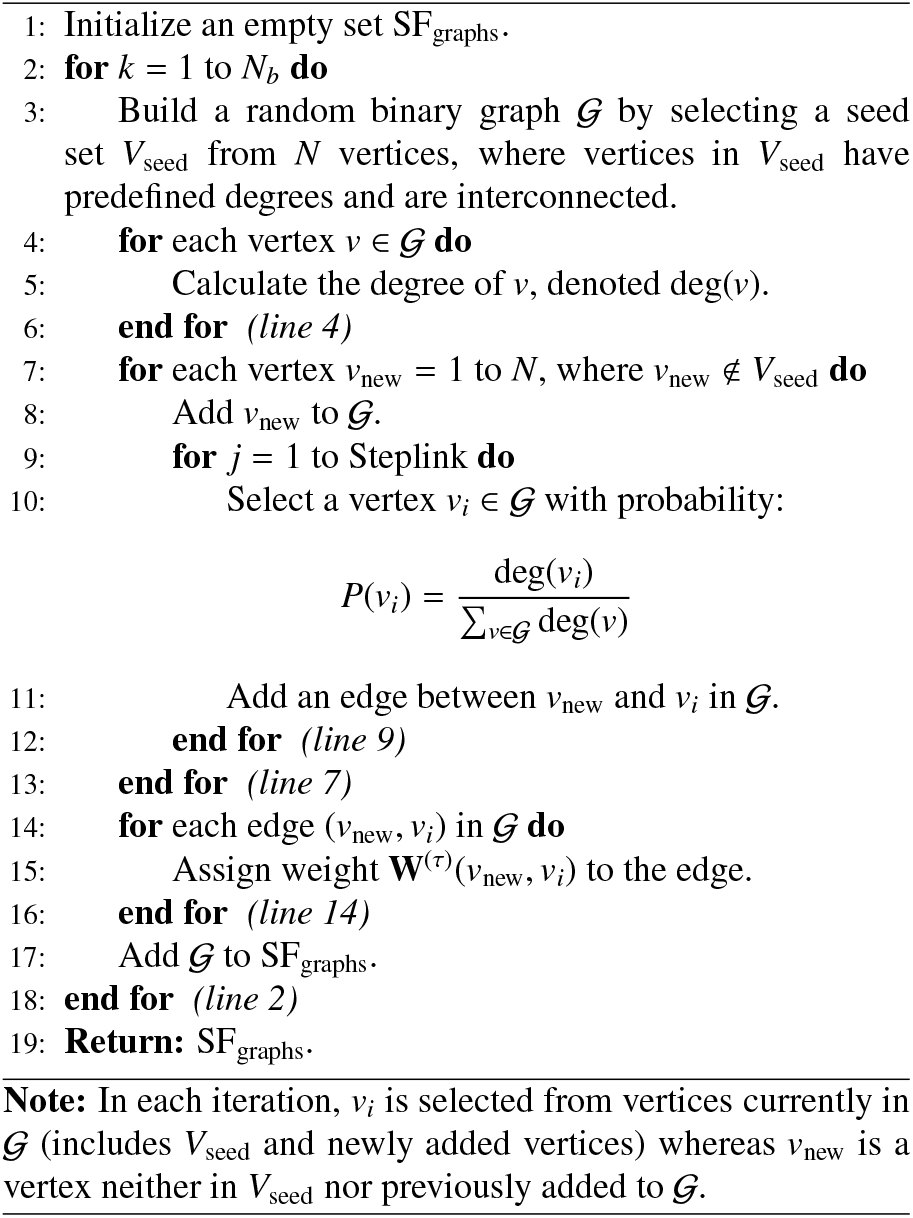

#### Algorithm 3

Small-World Scale-Free Graph Design

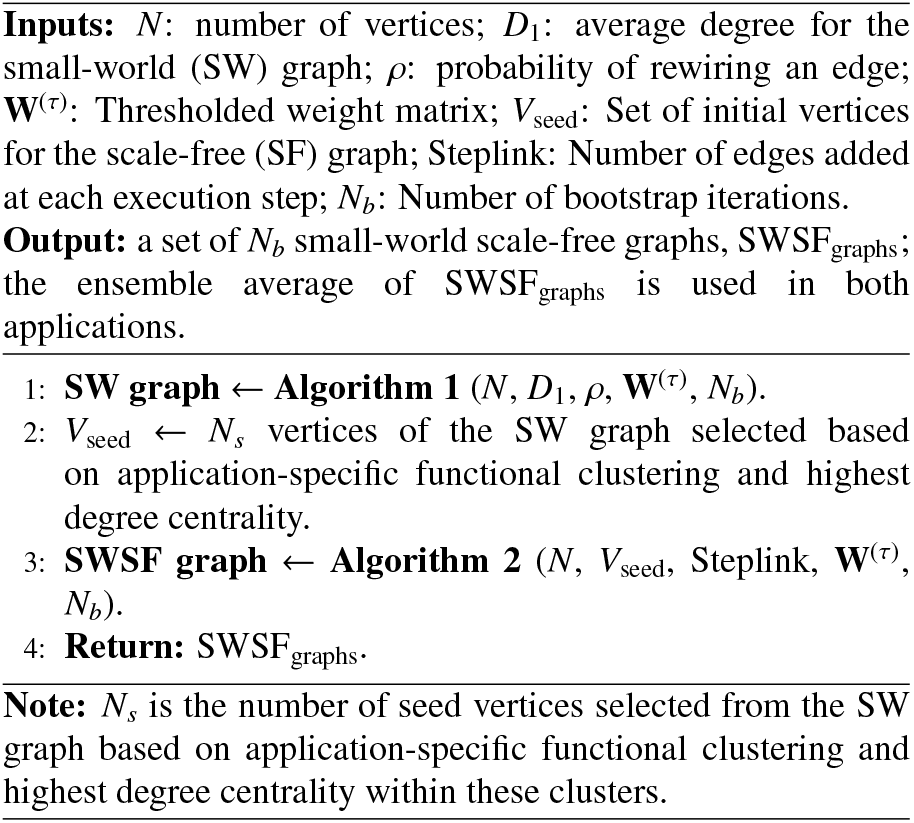

#### Determining the free parameters

Our method includes several free parameters, which were chosen either based on prior studies or optimised using cross-validation (CV) on the training set. CV was performed on Dataset-1 for the MI decoding task (see Fig. S1 in the Supplementary Material). For the brain fingerprinting application on Dataset-2, parameters from the MI decoding task were reused where applicable, and others optimised independently. A detailed 10-fold CV strategy—using classification accuracy as the selection criterion—ensured that our parameter choices were data-driven, robust, and reproducible. Full details of the CV procedure are provided in the Supplementary Material.

Selection of τ. The threshold parameter τ was determined through CV on Dataset-1 (MI decoding task), evaluating values from 0.2 to 0.8 using classification accuracy as the criterion (see Fig. S1, top left, in the Supplementary Material). The optimal value was determined to be τ = 0.4, which yielded graphs with an average degree of *D* = 29.40 ± 1.52. This value balances graph sparsity and connectivity, ensuring an accurate representation of the underlying network properties. The same threshold τ was applied to Dataset-1 (MI decoding) and Dataset-2 (brain fingerprinting) for a consistent graph construction.

Joint Selection of ρ and *D*. To enforce small-world properties, we restricted the rewiring probability (ρ) from the broader range 0.1 < ρ < 0.9 to 0.1 < ρ < 0.5, as values above ρ = 0.5 did not satisfy the small-world criteria: *I* > 1 and *P* > 0.6 (Fig. 2). This refinement, supported by prior studies [15, 17, 49], provided acceptable *I* and *P* values. Detailed results on graph metrics and classification performance for varying values of ρ and *D* are presented in Fig. S2 and Table S1 of the Supplementary Material.

**Fig. 2:**
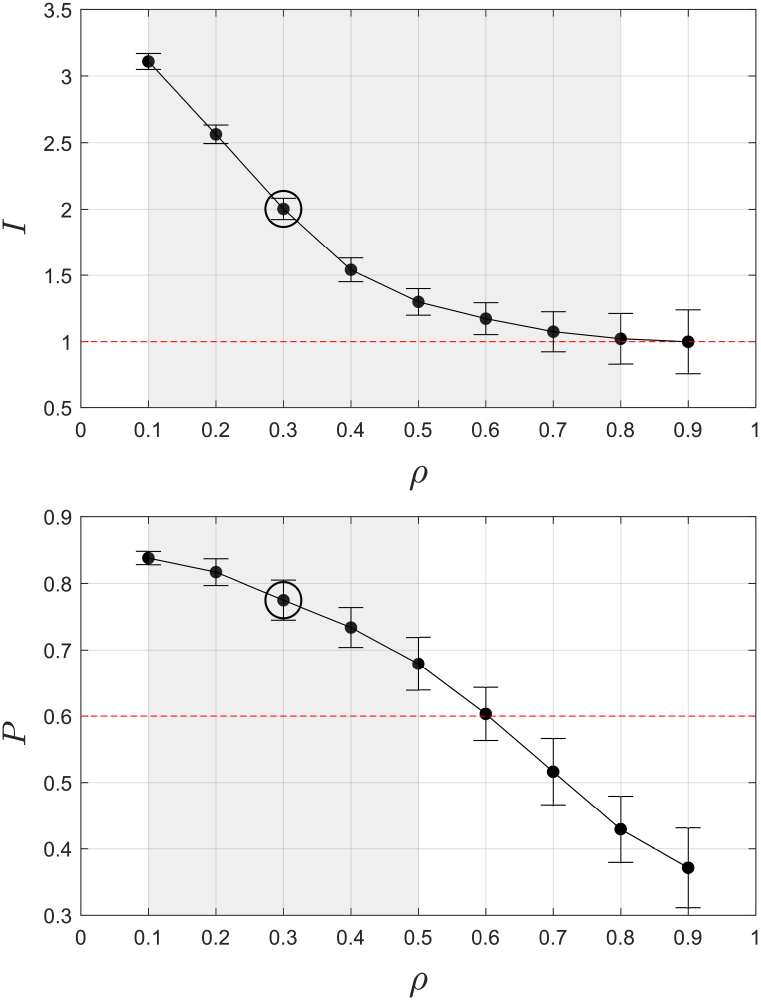
Effect of ρ on small-world measures *I* and *P*; value represent the mean across subjects. The gray area indicates the acceptable values for ρ, and the red dashed line is the threshold for the small-worldness of the graph. Error bars indicate the standard deviation across subjects and bootstraps.

A joint grid search was conducted over a two-dimensional parameter space: ρ (0.1 to 0.5, step 0.1) and average degree *D* (12 to 26, step 2), guided by prior studies [16, 17, 48] to balance structural properties and MI decoding accuracy. The optimal values—*D* = 20 (*Ē* = 0.17) and ρ = 0.3—achieved the highest training accuracy. Results are shown in Fig. S1 (top, right) in the Supplementary Material, highlighting small-world trade-offs.

Selection of *V*_seed_ nodes in SWSF graphs. To construct Small-world Scale-free (SWSF) brain graphs for MI decoding, we first created Small-world graphs with *N* = 118 nodes representing all EEG electrode regions. From these graphs, we selected 21 nodes (denoted as *V*_seed_) based on an application-specific functional clustering strategy as in previous related work [62]. Specifically, We grouped electrodes into functional clusters related to motor imagery and selected nodes with the highest degree centrality within each cluster, ensuring spatial relevance and structural importance. As central regions like the motor and somatosensory cortices are more effective for MI decoding, we selected a final set of 21 nodes proportionally across clusters to balance performance and computational efficiency. The choice of 21 nodes for *V*_seed_ is justified in Fig. S1 (bottom, left) of the Supplementary Material, showing its effect on MI classification accuracy. The SWSF graph was then built using all 118 nodes, with *V*_seed_ defining the scale-free substructure. This maintained a comparable edge count to the SW and SF graphs, enabling fair structural and performance comparisons.

For brain fingerprinting, we used the same identified electrodes as in the motor imagery task, given their efficiency in capturing activity patterns. Following earlier studies on brain fingerprinting [21, 63, 64], the frontal and parietal regions, known for their efficiency in this application, were prioritized during node selection. These regions guided the selection of 21 *V*_seed_ nodes used to construct the SWSF graph with *N* = 64 nodes for brain fingerprinting. The selected electrodes are visualized in Fig. 3, which presents the aggregate mean and standard deviation of their topographic distributions across subjects for each of the six frequency bands. The mean maps indicate how often each electrode was chosen as a *V*_seed_, revealing consistent spatial patterns that reflect robust functional organization. In contrast, the variability maps highlight inter-subject differences: higher standard deviations point to subject-specific roles, while lower values indicate electrodes that are consistently selected or consistently ignored. Individual subject topographies are shown in Fig. S3 of the Supplementary Material. This frequency-dependent variability aligns with established functional specializations across frequency bands.

**Fig. 3:**
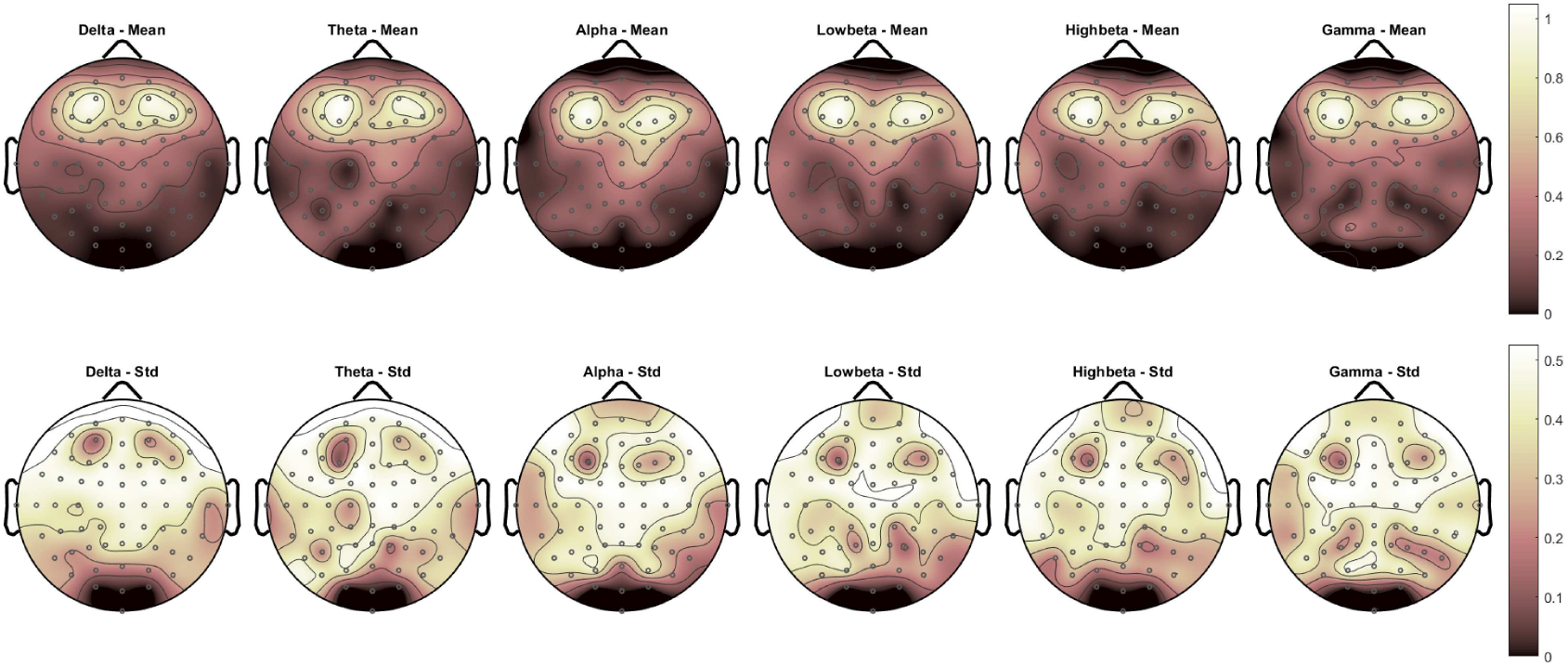
Mean (top) and standard deviation (bottom) topographic plots of the 21 selected vertices (*V*_seed_) across 109 subjects in the brain fingerprinting dataset, shown for all six frequency band.

*SF metrics*. One of the frequently used approaches to examine whether a network exhibits scale-free properties is analyzing its degree distribution. If the network follows a power-law, the number of nodes with degree *k*, denoted by *N*(*k*), should follow a distribution of the form *P*(*k*) ∼ *k*^−γ^, where the scaling exponent γ is typically in the range [2, 3]. In our study, we computed the histogram of node degrees and plotted the degree distribution on a log-log scale. In many cases, this yielded a linear trend in the tail of the distribution, with slope values (i.e., γ) within the expected range (Fig. S4). Although we refer to this behavior as power-law-like, it is more precise to state that the observed distributions are heavy-tailed, approaching power-law behavior asymptotically as *N*→∞. This aligns with the inherently finite nature of EEG graphs. Moreover, the coefficient of determination (*R*^2^) of the fitted line to the tail regions was computed to assess the goodness-of-fit, and in most instances, relatively high *R*^2^ values were obtained, indicating a reasonable fit to a heavy-tailed model. For each subject, we calculated the log-log plot, the *R*-squared value, and the slope of the regression curve to evaluate the adherence of SF graphs to a heavy-tailed degree distribution, following the methodology outlined in [16].

To support the scale-free characterization, we applied three Extreme Value Theory estimators—Hill, Moment, and Kernel—to the degree sequences of representative brain graphs. These provide statistically robust tail estimates, improving on naive slope-fitting methods. Figure S5 displays the empirical log-log CCDFs (black dots) overlaid with the fitted curves from each method across five subjects. Across subjects, Hill and Kernel estimators yield γ values in the range of approximately 2.5 to 3, consistent with classic scale-free regimes. In contrast, the Moment estimator returns larger values, roughly 3.5 to 5.5, likely due to its known upward bias in finite samples with noisy tails. These discrepancies highlight the importance of using multiple estimators to assess the robustness of tail exponent estimates.

To further interpret these exponents, we apply Voitalov’s taxonomy based on the tail index ξ = 1/(γ − 1). The estimated ξ values from the Hill and Kernel methods fall around 0.5 to 1, placing most graphs in either power-law with divergent second moment or power-law. The Moment estimator’s higher γ values imply ξ < 0.25, suggesting a few graphs fall into hardly power-law [65]. These results support the presence of heavy-tailed degree distributions across EEG-based brain graphs.

### 2.4. Datasets

This study employed two datasets: (i) motor-imagery task data from the BCI Competition III Dataset IVa [66, 67], and (ii) resting-state EEG data sourced from PhysioNet [68-70]. In the remainder of this paper, we refer to these two datasets as Dataset-1 and Dataset-2, respectively.

*Dataset-1*: Dataset IVa from BCI competition III consists of EEG signals from five healthy subjects, acquired using 118 electrodes at a sampling rate of 1000 Hz. A 2D projection of the electrode layout is shown in Fig. 4 (Left). Subjects were presented with visual cues for a duration of 3.5 seconds, during which they performed MI tasks involving their right hand or right foot. Random interruptions in the display of target cues, lasting between 1.75 and 2.25 seconds, served as rest periods for the subjects [66, 67]. We band-pass filtered the time-course of each electrode using a 5th-order Butterworth filter with a passband of 8-30 Hz to retain sensorimotor rhythms: mu (8-13 Hz) and beta (13-30 Hz), and subsequently down-sampled the signals to a frequency of 100 Hz. We then extracted, for each subject, 280 trials related to two classes of MI tasks (right hand and right foot), 140 trials per class. Each trial lasted 3.5 seconds (from 0.5 to 4 seconds after visual cue presentation), consistent with prior related work [71, 72].

**Fig. 4:**
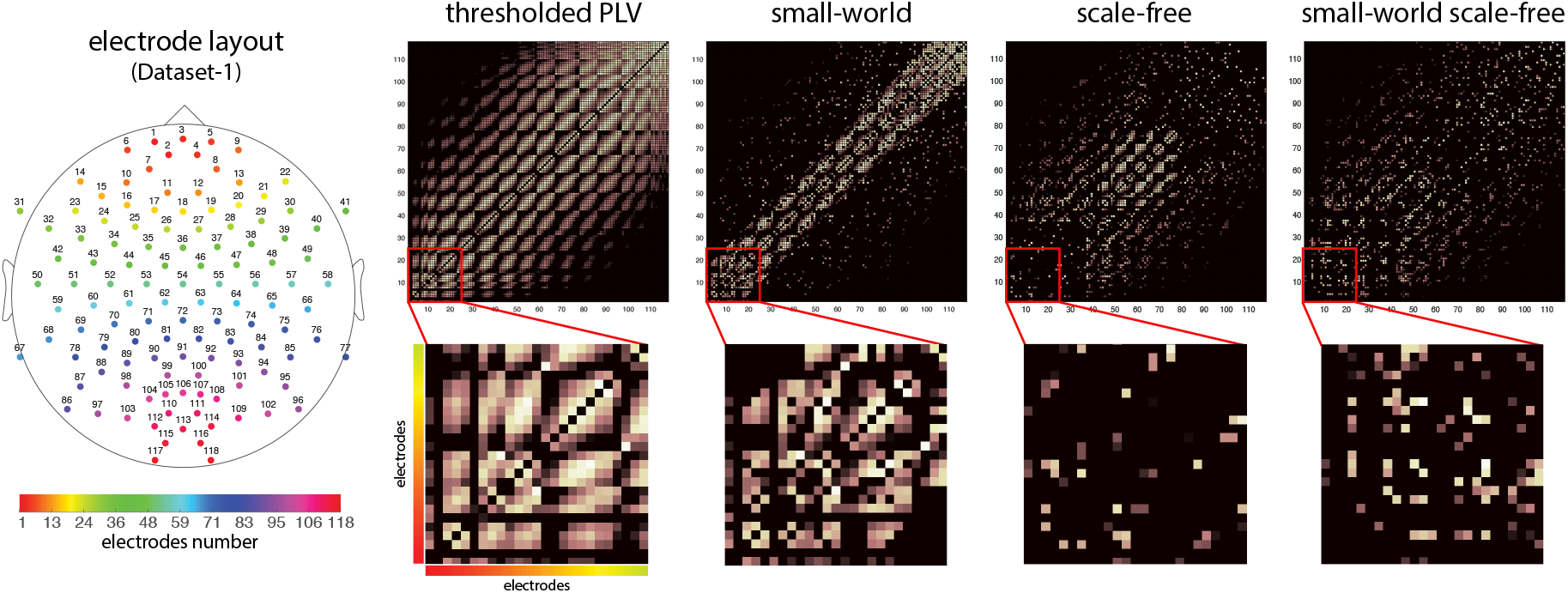
Adjacency matrices for the thresholded PLV, SW, SF, and SWSF graphs of subject aa; cortical location of electrodes are shown on the left; the correspondence between electrode positions and anatomical regions is shown in Fig. S6(b) in the supplementary material. Adjacency matrices for the PLV, SW, SF, and SWSF graphs of the other subjects of Dataset-1 are shown in Fig. S7 in the Supplementary Material, respectively.

*Dataset-2*: The resting-state EEG data from PhysioNet comprises EEG signals from 109 subjects, captured by 64 electrodes via the BCI2000 system [68, 69] at a sampling rate of 160 Hz. The location of 64 EEG electrodes in brain fingerprinting from Dataset-2 is shown in Fig. S6(a) in the supplementary material. Here, we used a subset of this dataset consisting of one-minute resting-state runs per subject (in the “eyes open” mode). For Dataset-2, we temporally bandpass filtered the resting-state EEG signals into six canonical frequency bands: delta (0.5-4 Hz), theta (4-8 Hz), alpha (8-13 Hz), low beta (13-20 Hz), high beta (20-30 Hz), and gamma (30-50 Hz).

### 2.5. Application 1: MI Decoding

To perform MI decoding, we used features derived either directly from the EEG signals or by using the inferred SW, SF, SWSF, and Kron-reduced EEG graphs. Firstly, we extracted a range of classical features from the EEG signals. These features fall into eight broad categories: statistical, Hjorth, variation, energy, entropy, band power, time series, and those based on common spatial patterns (CSP). We also extracted two GSP-based features named *DTV*(**f**_*i*_, **f** _*j*_) and **ē**_ℱ_. Given the large set of extracted features, we aimed to reduce their number to prevent redundancies, decrease computational cost, and improve performance generalization. Various methods have been proposed for feature selection within the context of MI decoding [73-76]. To this end, we used a meta-heuristic method known as differential evolution (DE) [77], a technique that has demonstrated superior performance in several related studies [62, 78]. We used the best subset of 10 features determined by this technique. These features were then used for classification using a support vector machine classifier with the radial basis function. We assessed the robustness of classification results using 10-fold cross-validation, with 10 rounds of repetition.

### 2.6. Application 2: Fingerprinting

For Dataset-2, we preprocessed the resting-state EEG signals recorded during the eyes-open mode by applying temporal bandpass filters to separate them into six standard frequency bands: delta (0.5-4 Hz), theta (4-8 Hz), alpha (8-13 Hz), low beta (13-20 Hz), high beta (20-30 Hz) and gamma (30-50 Hz). Data in each band was split into two parts, treating the initial 30 seconds as the test set and the second 30 seconds as the retest set. We then generated PLV, SWSF, and SWSF_kron_ brain graphs for both the test and retest sets for each of the 109 subjects. The feature sets were defined as the vectorized form of the upper triangular section of the adjacency matrices corresponding to these graphs.

Maximizing human functional connectome fingerprinting is based on the principle that two connectivity profiles derived from the same subject should exhibit greater similarity than two connectivity profiles derived from two different subjects. We assessed the fingerprinting power of our proposed graphs by deriving identifiability measures introduced in [22], as previously used in the context of EEG [63] and MEG [79] data. Specifically, given *S* subjects, we denote the test and retest feature vectors of subject *j* as 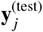 and 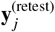 respectively, where *j* = 1, …, *S*. We then construct the *S* × *S* identifiability matrix whose (*i, k*) entry is the PLV value computed between 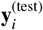 and 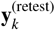. This matrix is not necessarily symmetric [79, 80].

Using the identifiability matrix (109 × 109 for each frequency band), a scalar quantity named *I*_self_ can be defined as the average of the main diagonal elements. These elements represent the PLV value between the test and retest feature sets of identical subjects, indicating the average self-similarity across subjects. Similarly, another scalar value named *I*_others_ can be defined as the average of the off-diagonal elements. These elements represent the PLV value between the test and retest feature sets of different subjects, quantifying the average cross-subject similarity based on the given feature set. Additionally, a third scalar quantity named *I*_diff_ = (*I*_self_ − *I*_others_) ×100, can be defined. This value provides a robust group-level estimate of identifiability at the individual connectome level, where a higher value indicates greater individual fingerprinting [22, 79].

## 3. Results and Discussion

This section presents an evaluation of the mentioned brain graph designs, MI decoding results derived from SWSF and SWSF_Kron_ brain graphs, along with brain fingerprinting results based on SWSF and SWSF_Kron_ brain graphs. Specifically, we analyze the influence of the accompaniment of GSP-based and classical features, the results of the Kron reduction efficacy in the proposed approach, and the accuracy of the SVM-RBF using various brain graphs to assess the separability of two-class MI tasks. Additionally, we discuss the fingerprinting capabilities of SWSF and SWSF_Kron_ over PLV-based graph and the influence of EEG duration on brain fingerprinting.

### 3.1. Graph-based Evaluation of Architectures

The adjacency matrix of the thresholded PLV, SW, SF, and SWSF graphs of a representative subject are shown in Fig. 4. In the SW structure, electrodes are predominantly interconnected with their neighbouring electrodes in both the clockwise and anti-clockwise directions. Consequently, the adjacency matrix of the SW brain graph exhibits a dominant banded structure around the main diagonal, reflecting the preserved local connectivity among spatially adjacent electrodes. In the SF graph, based on “preferential attachment,” the highest edge weights within the adjacency matrix are observed around the initial electrodes. Given the physiology of the motor cortex channels (used for MI decoding), we have placed these initial electrodes in the central brain regions. In the SWSF graph, high-significance nodes connect to adjacent ones, forming a SW structure visible as diagonal-adjacent stripes in the adjacency matrix. Increasing connections among these nodes introduces SF-like “preferential attachment,” evident in the matrix’s central rows and columns during MI decoding.

The combined use of PLV and small-world properties enables linking oscillatory phase synchrony to large-scale network topology. While PLV is sensitive to phase coupling that may reflect common sources, it remains a widely used and interpretable index of inter-regional coordination. Small-world metrics subsequently summarize how these connections organize into efficient, integrated networks, a framework well supported in network neuroscience.

Figure 5 illustrates the degree distribution of SWSF brain graphs, which demonstrates heavy-tailed characteristics, as opposed to Gaussian distribution traits. To examine the SWSF brain graphs more precisely, we computed both the SW and SF measurements for all subjects; see Table 2. The mean *C* and mean *L* of the SWSF graphs are 0.234 and 3.449, respectively, whereas the mean *I* and *P* values are 1.041 and 0.640, respectively, which reflect the small-worldness of these graphs. The same measures but for the SW graphs are presented in Table S2 in the Supplementary Material. It is worth noting that scale-free and small-world features may coexist in brain networks. Hubs characteristic of scale-free organization can promote small-world efficiency by reducing path lengths, but the emergence of small-world topology does not inherently modify the heavy-tailed form of scale-free degree distribution.

**Table 2:** Graph metrics for the constructed SWSF brain graphs in the MI decoding application. For all graphs and subjects: *N* = 118.

| Metric | aa | al | av | aw | ay | Average |
| --- | --- | --- | --- | --- | --- | --- |
| $C$ | 0.242 | 0.226 | 0.222 | 0.238 | 0.243 | 0.234 |
| $L$ | 3.422 | 3.460 | 3.447 | 3.431 | 3.485 | 3.449 |
| $C/C^{(rnd)}$ | 1.502 | 1.466 | 1.471 | 1.487 | 1.519 | 1.489 |
| $L/L^{(rnd)}$ | 1.417 | 1.437 | 1.430 | 1.425 | 1.442 | 1.430 |
| $I$ | 1.060 | 1.020 | 1.029 | 1.044 | 1.053 | 1.041 |
| $\Delta C$ | 0.506 | 0.485 | 0.492 | 0.513 | 0.477 | 0.495 |
| $\Delta L$ | 0.121 | 0.092 | 0.116 | 0.108 | 0.147 | 0.117 |
| $P$ | 0.632 | 0.651 | 0.643 | 0.629 | 0.647 | 0.640 |
| $\gamma$ | 2.51 | 2.45 | 2.40 | 2.36 | 2.34 | 2.41 |
| $R^2$ | 0.84 | 0.85 | 0.85 | 0.84 | 0.86 | 0.85 |
| $N_h$ | 23 | 22 | 22 | 22 | 21 | 22 |

**Fig. 5:**
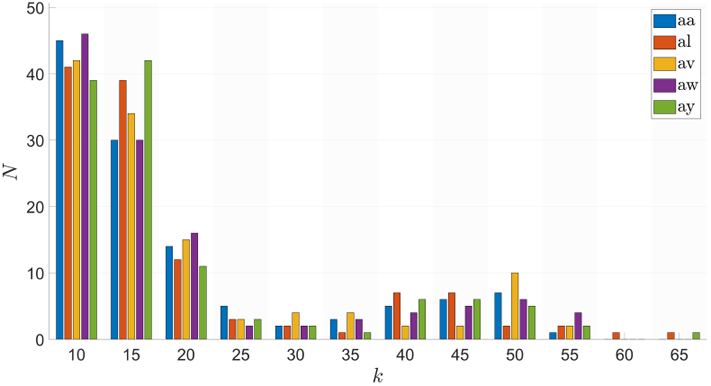
Degree distributions of the SWSF graphs for all five subjects in the MI decoding application.

Furthermore, we found that the average exponent of the fitted power-law-like distribution for subjects in the SWSF design is approximately 2.41, with an *R*-squared value close to 0.85. Therefore, in comparison with the results obtained for the SF measures, it becomes evident that the application of the SW scheme leads to a reduction in both the exponent parameter and *R*-squared, which reflects the scale-free nature of these networks. The average number of hubs (*N*_*h*_) for the SWSF graph across subjects is 22, nearly identical to SF graphs.

The SW_Kron_, SF_Kron_, and SWSF_Kron_ graphs for subjects aa, al, av, aw, and ay are each comprised of 24 selected nodes and 276 corresponding edges, see Fig. 6. The three Kron-reduced graphs (SW_Kron_, SF_Kron_, and SWSF_Kron_) originate from the same PLV-based graph, which consists of 118 nodes. Subsequently, these graphs are derived from the SW, SF, and SWSF graphs, each consisting of 118 nodes and characterized by distinct metrics and topologies.

**Fig. 6:**
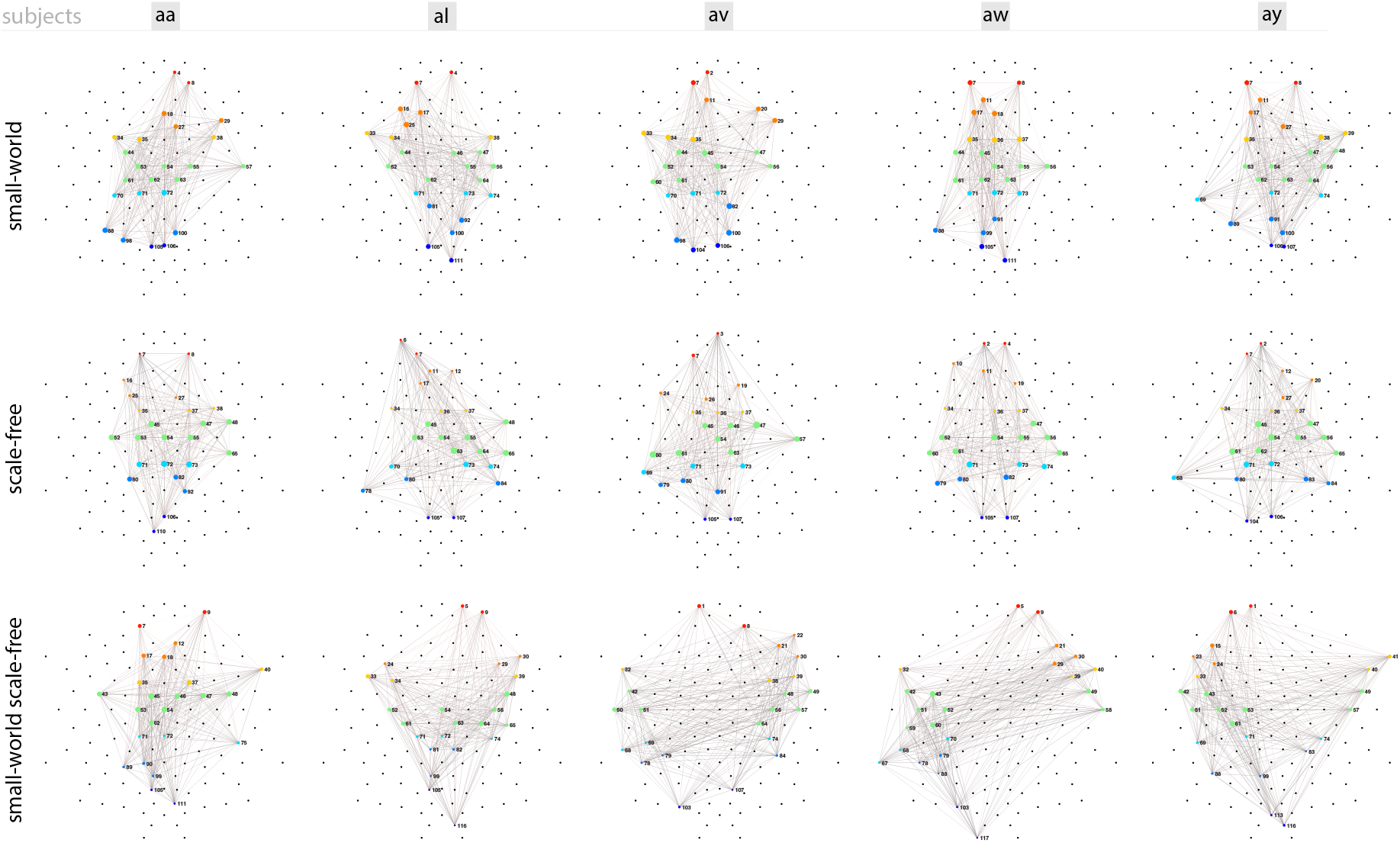
Kron-reduced SW, SF, and SWSF graphs across all five subjects in the MI decoding application.

### 3.2. Application 1: MI Decoding via SWSF Brain Graphs

MI decoding analysis was performed on Dataset-1. From the available set of classical and GSP features, the following top ten features from the different classes were selected through differential evolution (cf. Section 2.5), which consist of, GSP features: **ē**_ℱ_, *DTV*(**f**_*i*_, **f** _*j*_); temporal features: second difference, standard deviation, log root sum of sequential variations, mean Teager energy, log energy entropy; CSP features: CSP (1st filter); frequency features: *µ* band power; time series features: AR (4th coefficient). Note that the dimensionality of each feature type is as follows: the graph spectral feature **ē**_ℱ_ has dimensionality equal to the number of EEG channels per trial; the *DTV* feature has dimensionality equal to the number of time samples per trial; the time and time-series features also have dimensionality equal to the number of channels per trial; and the CSP-based features have dimensionality equal to the number of spatial filters retained per trial. The GSP and classical features are described in detail in Section 2.1 and Table S3, respectively. The same 10 features were selected across all graph designs. Based on DE feature selection, the best fitness function was obtained when GSP-based features were combined with classic MI features. Combining **ē**_ℱ_ and *DTV*(**f**_*i*_, **f** _*j*_) with classical features, including multiple time-domain features, the CSP feature, *µ* band power, and autoregressive (AR) coefficient, led to optimal results. We used a statistical two-sample t-test with a significance level of 0.01, to examine the level of distinguishability of the features. The t-test results showed that the variance ratio for the first CSP filter has the highest separability. Compared to the CSP feature, the GSP-based, time, and frequency features had lower absolute t-values and higher p-values.

Table 3 compares MI decoding classification accuracy results using five different methods: CLA, PLV, SW, SF, and SWSF. The SW, SF, and SWSF graphs outperform the PLV graph and CLA; the CLA method only uses classical features whereas PLV, SW, SF, and SWSF use both classical and GSP features as they are graph-based. The performance gradually increases when transitioning from the CLA model to the SWSF model, with SWSF showing superior performance over all the other methods. We also tested the results against a lightweight deep learning benchmark, specifically EEGNet, which is a compact convolutional neural network that integrates established EEG feature extraction principles for BCI applications [81]. We used a standard configuration for EEGNet consistent with previous motor imagery studies (e.g. [82]), with minor adjustments for regularization and optimization; specifically, the network used 8 temporal filters (F1 = 8), 2 depth multiplier (spatial filters) D = 2, 16 point-wise filters (F2 = 16), kernel length 64, dropout probability 0.5, norm rate 0.5, batch size 16, and a learning rate of 5 ×10^−4^.

**Table 3:** MI decoding accuracy (mean ± std,%) of the proposed method for six different models of brain graph (*N* = 118, *M* = 1180) using within-subject data. For CLA, the top 10 classical features were used whereas for PLV, SW, SF, and SWSF the top 10 GSP-based and classical features were used. Moreover, the results of EEGNet are shown as a representative lightweight deep learning architecture. The symbol * indicates that the SWSF model is significantly better than all other models, paired t-test, Bonferroni-corrected, *α*_*ad j*_ = 0.00167, p < 0.00167).

| Subject | CLA | PLV | SW | SF | SWSF | EEGNet |
| --- | --- | --- | --- | --- | --- | --- |
| aa | 78.93±3.00 | 79.64±3.21 | 82.14±2.30 | 80.36±2.71 | **84.29±1.73\*** | 68.93±4.14 |
| al | 92.14±3.19 | 90.00±3.13 | 95.00±1.95 | 92.86±2.16 | **96.07±1.60\*** | 93.21±2.64 |
| av | 72.86±2.54 | 73.21±4.38 | 78.93±2.61 | 73.93±2.81 | **79.29±2.22\*** | 60.36±3.55 |
| aw | 82.14±2.77 | 83.93±3.87 | 88.57±2.45 | 85.36±2.02 | **90.71±1.93\*** | 72.50±1.73 |
| ay | 83.21±2.48 | 82.14±4.55 | 88.57±2.63 | 82.86±2.66 | **91.79±1.72\*** | 69.29±1.84 |
| mean±std | 81.86±7.02 | 81.78±6.13 | 86.64±6.27 | 83.07±6.93 | **88.43±6.62\*** | 72.86±12.24 |

The results of MI decoding using the Kron-reduced graphs are reported in Table 4. Kron reduction enhanced the classification accuracy across all graphs. The SWSF_Kron_ model outperforms SW_Kron_, SF_Kron_, and PLV_Kron_ models in all subjects. Moreover, a comparison of Tables 3 and 4 shows that the average performance of all four models improved when combined with Kron reduction, demonstrating the utility of this method. Compared to the original schemes (PLV, SW, SF, and SWSF), the average MI decoding accuracy in the Kron-reduced schemes (PLV_Kron_, SW_Kron_, SF_Kron_, and SWSF_Kron_) increased by 1.65%, 2.79%, 1.79%, and 3.00%, respectively. As confirmed by the results in Table 4, similar to those in Table 3, SWSF brain graphs yield superior results in MI decoding compared to the SW and SF brain graphs.

**Table 4:** MI decoding accuracy (mean ± std,%) of the proposed method for four different models of brain graph with Kron reduction using within-subject data. The symbol * indicates that the SWSF_Kron_ model is significantly better than all other models (paired t-test, Bonferroni-corrected, *α*_*ad j*_ = 0.00167, p < 0.00167).

| Subject | PLV<sub>Kron</sub> | SW<sub>Kron</sub> | SF<sub>Kron</sub> | SWSF<sub>Kron</sub> |
| --- | --- | --- | --- | --- |
| aa | 80.71±5.88 | 85.36±2.12 | 81.79±2.03 | **88.21±1.50\*** |
| al | 92.50±3.05 | 97.14±2.26 | 93.57±1.76 | **98.57±1.94\*** |
| av | 75.00±3.13 | 80.36±1.99 | 76.07±2.05 | **82.86±1.97\*** |
| aw | 85.71±2.43 | 91.79±1.83 | 86.43±1.87 | **94.64±1.52\*** |
| ay | 83.22±4.99 | 92.50±2.16 | 86.43±1.68 | **92.86±1.48\*** |
| mean±std | 83.43±6.44 | 89.43±6.58 | 84.86±6.47 | **91.43±6.06\*** |

To assess the statistical robustness of performance differences in Dataset-1, we conducted paired t-tests on the 10-fold CV accuracies for each subject. After Bonferroni correction for multiple comparisons (*further contextualize the performance*_*adj*_ = 0.00167), SWSF and SWSF_Kron_ significantly outperformed all baseline graph models across all five subjects (p < 0.00167); see Tables 3 and 4. For completeness, detailed paired t-test results are presented in Tables S4 and S5 in the Supplementary Material.

Our main experiments have been focused on within-subject decoding, consistent with the goal of building individualized brain graphs that depend on each participant’s PLV structure and topological organization. However, we additionally conducted a cross-subject analysis on Dataset-1. Specifically, we adopted a strict leave-one-subject-out paradigm. In each fold, the model was trained using data from all subjects except the target subject, which was held out entirely for testing. This protocol evaluates the ability of each method to generalize to previously unseen individuals and provides a realistic assessment of subject-independent performance. The results revealed a reduction in accuracies compared to the within-subject setting, which is intuitive and was expected given the degree of inter-individual variability; see Tables S6 and S7 in the Supplementary Material. In particular, SWSF remained the top-performing model (Table S6), outperforming PLV by 8-14%, SW by 4-7%, and SF by 6-10%, and furthermore, SWSF_Kron_ remained the top-performing model (Table S7), outperforming PLV_Kron_ by 9-14%, SW_Kron_ by 3-8%, and SF_Kron_ by 6-12%. These results indicate that, despite being optimized for personalized modeling, the proposed method still exhibits competitive generalization to unseen subjects.

To further contextualize the performance of the proposed SWSF_Kron_ model, we expanded our evaluation to include several recent and representative GSP-based (GSP-FKT [72], GSL [83]) and CSP-based (*ℓ*_1_-CSP [84], RCCSP [85], BECSP [86]) approaches that have demonstrated strong performance in motor imagery decoding (Table 5).

**Table 5:**
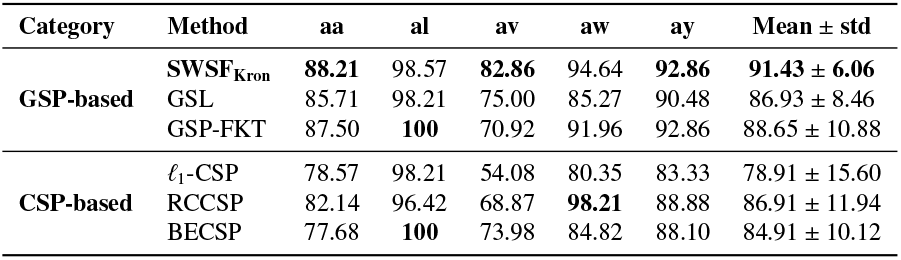
MI decoding accuracy (%) comparison between GSP-based and CSP-based methods. The best result in each column is boldfaced.

### 3.3. Application 2: Fingerprinting via SWSF Brain Graphs

Brain fingerprinting analysis was performed on Dataset-2. In the following, we compare the performance of PLV, SWSF, and SWSF_kron_ brain graphs in brain fingerprinting under varying rewiring probabilities and EEG time lengths.

Figure 7 shows fingerprinting results as a function of the rewiring probability for SWSF, and SWSF_kron_ graphs, across six frequency bands; the parameter *ρ* does not apply to PLV graphs. Optimal subject identifiability was found for ρ = 0.6 for the Delta and Theta bands, and *ρ* = 0.3 for the Alpha, Low-Beta, High-Beta, and Gamma bands. In both SWSF and SWSF_kron_ graphs, the Alpha and Beta bands, which are associated with both motor imagery and fingerprinting, exhibited the best performance for *ρ* = 0.3. In contrast, for the Delta and Theta bands — which we study only in the context of fingerprinting — the best identifiability was obtained for *ρ* = 0.6. Lastly, SWSF_kron_ generally outperforms SWSF across all six frequency bands, except for the Gamma band, where both methods demonstrated comparable performance.

**Fig. 7:**
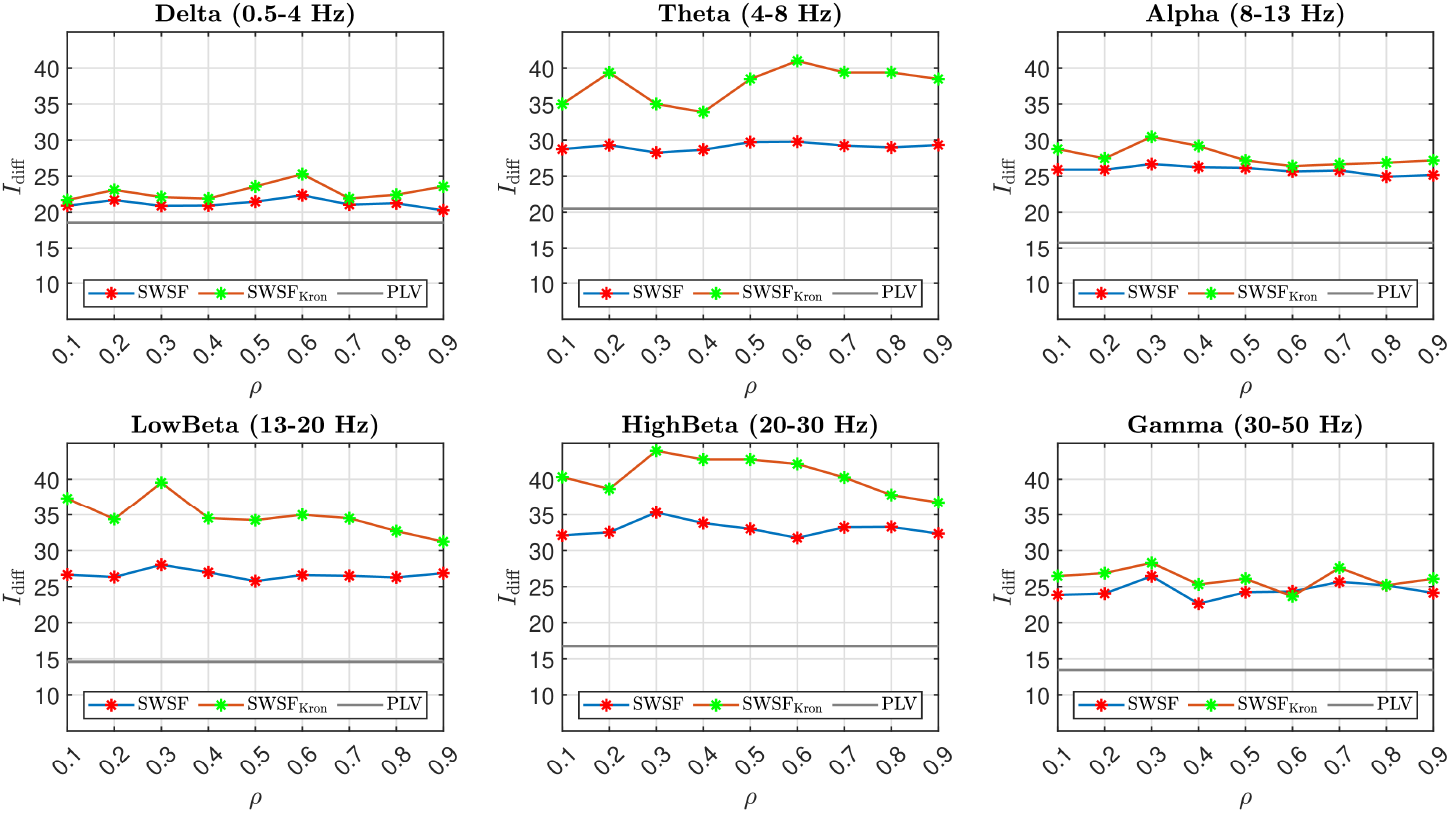
Differential identifiability power of PLV, SWSF, and SWSF_kron_ graphs as a function of the rewiring probability (ρ) across six frequency bands on Dataset-2.

Table 6 presents a comparison of the best performances of SWSF and SWSF_kron_ in brain fingerprinting across six frequency bands for the optimal *ρ*. As shown in Fig. 7, these results are benchmarked against PLV for evaluation. The results demonstrate a clear superior performance of the inferred graphs (SWSF and SWSF_kron_) in the *I*_diff_ measure, compared to the conventional PLV-based graphs, across all frequency bands.

**Table 6:** *I*_diff_ (%) for fingerprinting on Dataset-2. Statistical significance was assessed using the Wilcoxon signed-rank test across 109 subjects (Bonferroni-corrected *α*_*ad j*_ = 5.6 ×10^−4^). The symbol † indicates that the SWSF or SWSF_Kron_ model significantly outperformed PLV; the symbol ‡ indicates that SWSF_Kron_ significantly outperformed SWSF.

| Frequency-Band | PLV | SWSF | SWSF<sub>Kron</sub> |
| --- | --- | --- | --- |
| Delta | 18.56 | 22.35† | **25.28†** |
| Theta | 20.47 | 29.77† | **40.99$\dagger\ddagger$** |
| Alpha | 15.74 | 26.66† | **30.47$\dagger\ddagger$** |
| Low-Beta | 14.58 | 28.02† | **39.53$\dagger\ddagger$** |
| High-Beta | 16.75 | 35.30† | **43.96$\dagger\ddagger$** |
| Gamma | 13.44 | 26.45† | **28.29†** |

To investigate the impact of EEG time length on fingerprinting, we performed identification using varying trial lengths in Dataset-2. To enhance the stability and robustness of the investigation, the bootstrap method was employed to select random data subsets, with 30 repetitions. In Dataset-2, the length of EEG epochs used to derive the graphs was varied from 2 to 30 seconds (2, 3, 4, 5, 6, 7, 8, 10, 12, 15, 20, 30) for both the test (session 1) and retest (session 2) sets, due to limited trial availability. The power of identifiability is depicted as a function of the epoch length, ranging from 2 to 30 seconds, for all six frequency bands. Differential identifiability was calculated as the average of 30 bootstrap iterations for each trial instance. In each iteration, n trials (n seconds) were randomly selected from the EEG data of each subject across all six frequency bands, and the PLV, SWSF, and SWSF_kron_ brain graphs were subsequently derived.

Figure 8 shows fingerprinting performance versus resting-state data length. Overall, performance improves with longer recordings across all models, except in the Gamma and Theta bands. In the Gamma band, all models peak at 3 seconds, then show minor fluctuations up to 30 seconds. In the Theta band, the SWSFkron graph peaks at 3 seconds and slightly declines thereafter. As EEG length increases, *I*diff rises and its standard deviation decreases across all bands, enhancing identifiability and fingerprinting effectiveness.

**Fig. 8:**
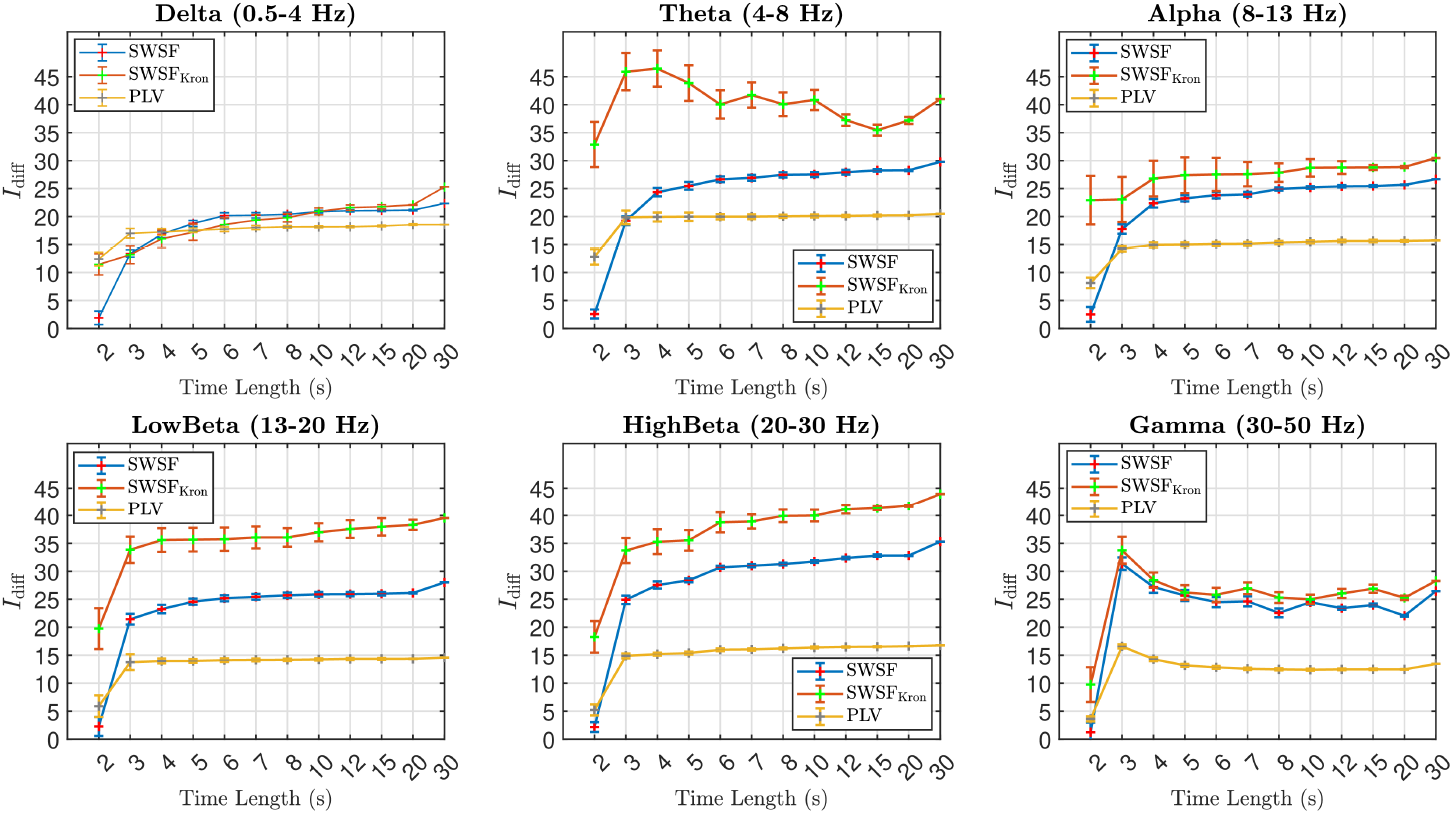
Differential identifiability power of PLV, SWSF, and SWSF_kron_ graphs as a function of the length of epochs used to derive the graphs for six frequency bands on Dataset-2. The error bars show the standard deviation across 30 bootstrap iterations. Statistical significance between models is assessed at the final 30-second window (Table 6), using Wilcoxon signed-rank tests across 109 subjects with Bonferroni correction (*α*_*ad j*_ = 5.6 × 10^−4^).

The observed fluctuations and occasional decline in fingerprinting performance at longer epoch lengths likely reflect the non-stationary nature of EEG signals. As longer temporal windows encompass more transient neural dynamics and noise, phase-based connectivity estimates such as PLV become less stable, reducing within-subject consistency. Moreover, longer segments reduce the number of independent epochs available for averaging, which can increase variance in identifiability metrics.

As shown in Fig. 8, the identifiability curves gradually increase and stabilize after approximately 6 to 8seconds, with the highest and most reliable performance obtained at the 30-second window. Following standard practice in EEG/MEG fingerprinting studies, statistical comparisons between PLV, SWSF, and SWSF_Kron_ were therefore conducted at this final stable window. Wilcoxon signed-rank tests across 109 subjects, with Bonferroni correction for 18 pairwise comparisons (*α*_*ad j*_ = 5.6 × 10^−4^), showed that both SWSF and SWSF_Kron_ achieved significantly higher *I*_diff_ values than PLV in all six frequency bands (p < 5.6 × 10^−4^) (Table 6). Moreover, SWSF_Kron_ significantly outperformed SWSF in the Theta, Alpha, Low-Beta, and High-Beta bands (p < 5.6 × 10^−4^). These results confirmed that the proposed graph models yield robust and statistically significant improvements in cross-subject EEG identifiability.

Across all settings, SWSF and SWSF_kron_ models consistently outperform PLV graphs, except for the Delta band where the performances are nearly identical. The most significant performance difference between SWSF and SWSF_kron_, in comparison to PLV graphs, is observed in the Low Beta and High Beta bands. A pattern that remains consistent across varying epoch lengths.

## 4. Conclusion

We proposed a novel method for constructing brain graphs from EEG data, generating subject-specific networks that reflect small-world and scale-free properties based on the phase-locking value between electrode time courses. To reduce complexity while retaining essential information, we applied Kron reduction. By combining graph-derived features with classical ones, we demonstrated improved performance in motor imagery decoding and brain fingerprinting across two independent datasets. These results underscore the potential of our graph design for broader EEG analysis and applications in cognitive neuroscience.

## CRediT authorship contribution statement

**Mohammad Davood Khalili:** Conceptualization, Data curation, Formal analysis, Investigation, Methodology, Software, Visualization, Writing—original draft, Writing—review & editing. **Vahid Abootalebi:** Conceptualization, Methodology, Supervision, Writing—review & editing. **Hamid Saeedi-Sourck:** Supervision, Writing—review & editing. **Andrea Santoro:** Methodology, Writing—review & editing. **Harry H. Behjat:** Conceptualization, Investigation, Methodology, Software, Supervision, Visualization, Writing—review & editing.

## Ethics Statement

This research study was conducted retrospectively using human subject data made available in open access. Ethical approval for processing the data was not required as confirmed by the license attached with the open access data. The data analyzes that we performed are in compliance with relevant local laws and institutional guidelines.

## Data and Code Availability

The datasets used in this paper are publicly available; Dataset-1: www.bbci.de/competition/iii/; Dataset-2: https://physionet.org/content/eegmmidb/1.0.0/.

We customized four MATLAB toolboxes to implement various stages of our proposed approach. Specifically, EEGLAB [87] for EEG preprocessing and scalp projection, Brainstorm [88] for brain projection, GSPBOX [89] for generating the graph figures, and the Brain Connectivity Toolbox [90] for calculating graph metrics.

## Acknowledgments

Dataset-1 used in this work was provided as part of the BCI Competition III, organized by the Berlin Brain-Computer Interface team (https://www.bbci.de/). Dataset-2 was made available by the developers of the BCI2000 instrumentation system for brain-computer interface research (https://www.bci2000.org/downloads/doc/paper.pdf).

H.H.B. received funding from the Swedish Research Council under grant no. 2018-06689 and the European Union, Horizon Europe Research and Innovation Program under Marie Skłodowska-Curie action under grant no. 101153323. A.S. acknowledges funding from the European Union, Horizon Europe Research and Innovation Program under Marie Skłodowska-Curie action under grant no. 101208090

---

**Fig. S1:**
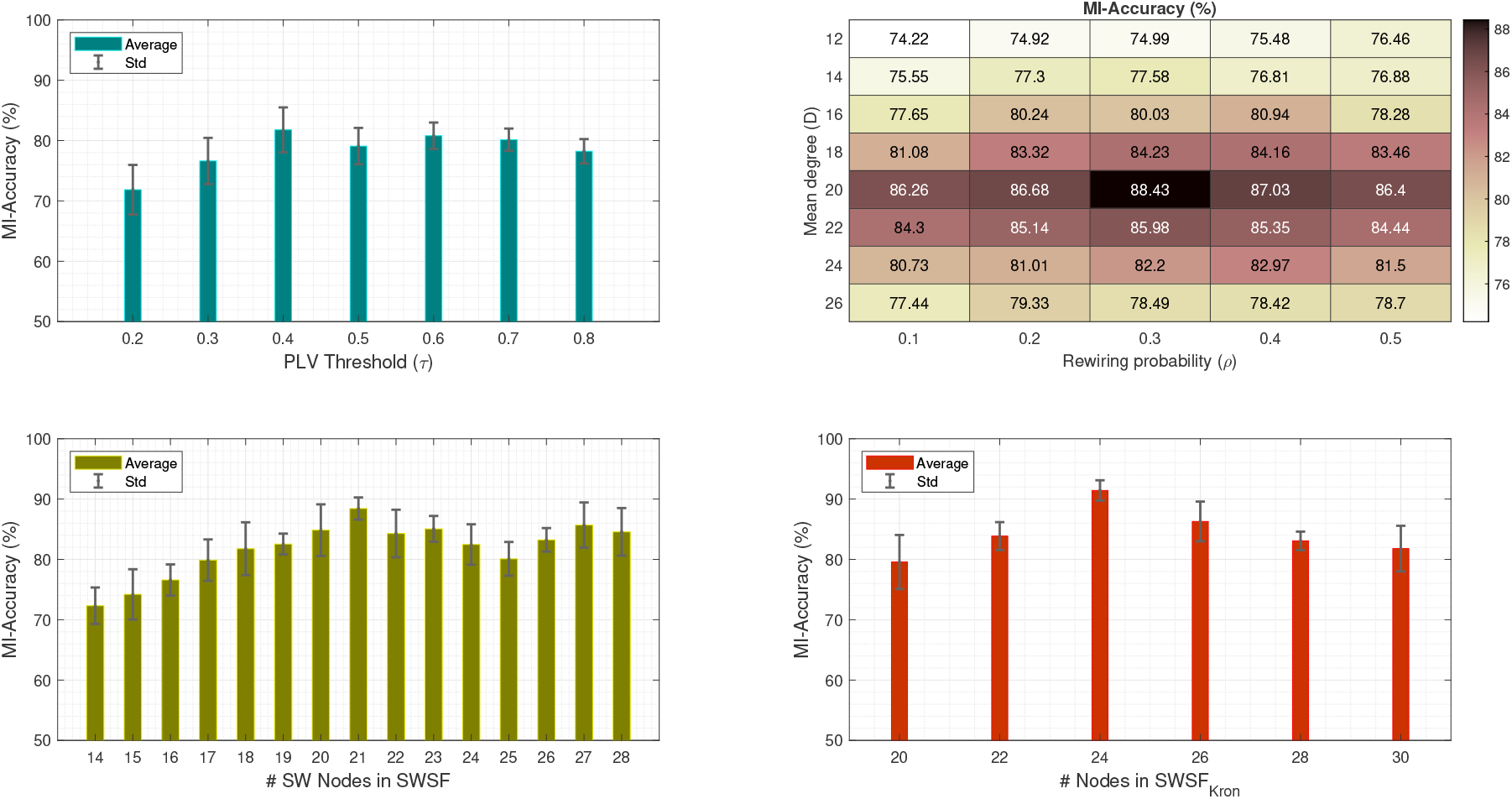
Cross-validation strategy for parameter selection. To ensure generalization across tasks, a systematic 10-fold cross-validation (CV) was applied to both motor imagery (MI) decoding (Dataset-1) and brain fingerprinting (Dataset-2). Five parameters were optimized: PLV threshold (*τ*), rewiring probability (*ρ*), average degree (*D*), initial seed vertices (*V*_seed_), and selected vertices for Kron reduction. For MI decoding, *τ* was tuned independently, while *ρ* and *D* were optimized jointly via grid search. *V*_seed_ and Kron-reduction nodes were selected using a hybrid strategy combining functional clustering with degree centrality, followed by empirical evaluation to identify subsets maximizing validation accuracy. (Top-left) shows performance across *τ* values; (Top-right) shows joint tuning of *ρ* and 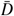. (Bottom-left) explores varying the number of *V*_seed_ nodes in the SWSF graph; (Bottom-right) shows accuracy trends for different node selections in the Kron-reduced graphs. For brain fingerprinting, *ρ* and *D* were re-optimized using CV, while *τ, V*_seed_, and Kron nodes were transferred from the MI task. Biological plausibility constraints (*I* > 1, *P* > 0.6) were enforced to ensure small-world structure.

**Table S1:** Classification accuracy in MI decoding using small-world brain graphs derived by using different values of the free parameter *ρ*.

| Subjects | SW($\rho = 0.1$) | SW($\rho = 0.2$) | SW($\rho = 0.3$) | SW ($\rho = 0.4$) | SW($\rho = 0.5$) |
| --- | --- | --- | --- | --- | --- |
| aa | 74.29±1.89 | 78.57±2.64 | 82.14±2.43 | 80.36±2.26 | 75.00±2.05 |
| al | 83.93±1.94 | 89.64±1.44 | 95.00±2.17 | 91.79±2.54 | 84.64±1.89 |
| av | 75.71±2.17 | 77.86±2.21 | 78.93±2.29 | 76.43±2.34 | 75.36±2.41 |
| aw | 78.21±2.18 | 82.14±2.57 | 88.57±2.15 | 84.29±2.27 | 81.79±1.84 |
| ay | 76.43±2.26 | 81.43±2.08 | 88.57±1.99 | 83.92±2.39 | 77.86±2.28 |
| Mean±std | 77.71±2.09 | 81.93±2.19 | 86.64±2.21 | 83.36±2.36 | 78.93±2.09 |

**Table S2:** Graph metric values used to validate the small-world characteristics of the constructed small-world brain graphs. For all graphs and subjects: *N* = 118, *M* = 1180, *D* = 20, *Ē* = 0.17, and *ρ* = 0.3.

| Metric | aa | al | av | aw | ay | Average |
| --- | --- | --- | --- | --- | --- | --- |
| $C$ | 0.361 | 0.384 | 0.369 | 0.372 | 0.377 | 0.373 |
| $L$ | 2.088 | 2.145 | 2.072 | 2.168 | 2.123 | 2.119 |
| $I$ | 2.047 | 1.963 | 2.028 | 2.008 | 1.969 | 2.003 |
| $P$ | 0.785 | 0.769 | 0.781 | 0.776 | 0.764 | 0.775 |

**Fig. S2:**
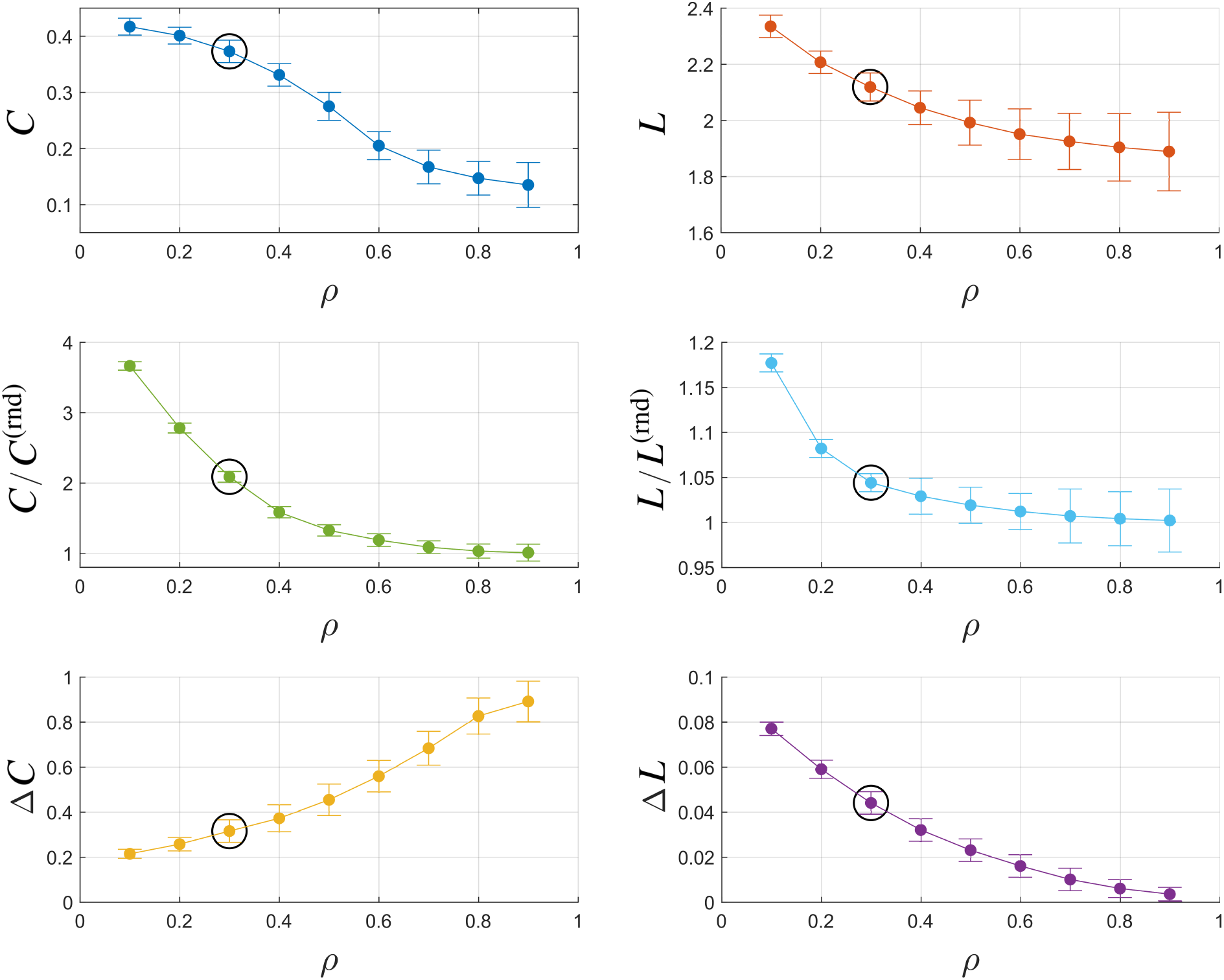
Effect of *ρ* on *C, L, C/C*^(rnd)^, *L/L*^(rnd)^, Δ*C*, and Δ*L* for SW brain graphs over all subjects in MI decoding.

**Fig. S3:**
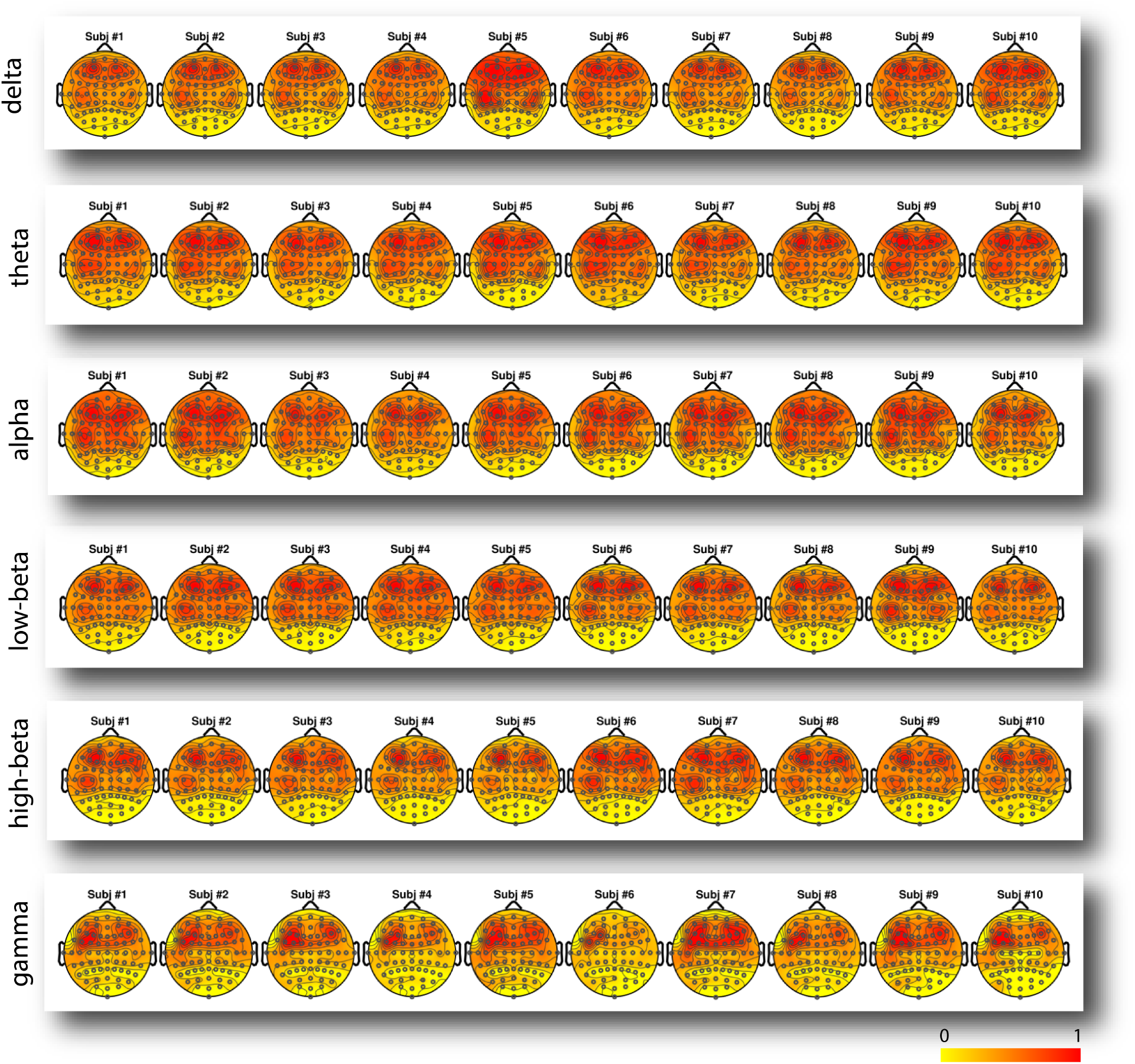
Topographic plots showing electrode significance for brain fingerprinting across the first 10 subjects of Dataset-2, displayed by frequency band.

**Fig. S4.**
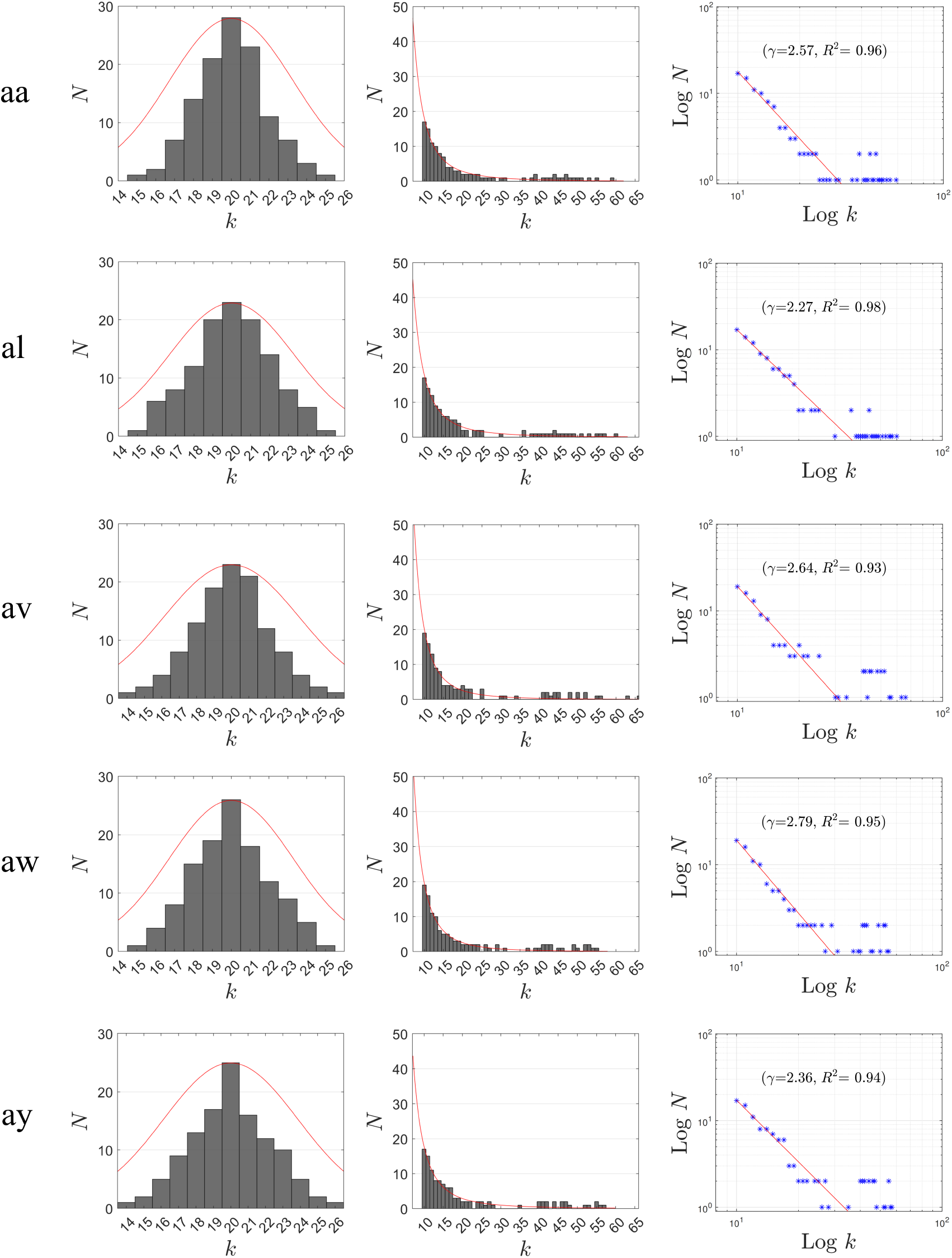
Left: Histogram of small-world degree distribution with their corresponding fitted Normal distribution. Center: Histogram of scale-free degree distribution (center) with fitted power-law curves, exhibiting exponents between 2 and 3. Right: log-log plots highlighting scale-free properties, with corresponding *γ* values and *R*^2^ scores shown. The number of hubs (*N*_*h*_) is 21 for aa, al, av, aw and 20 for ay, based on 118 electrodes, 1180 edges, and *D* = 20. The average power-law exponent across subjects is 2.53 with *R*^2^ ≈ 0.95. Hubs are defined as nodes with degrees at least 1.5 times the mean, averaging 21 hubs (18% of nodes) per subject.

**Table S3:**
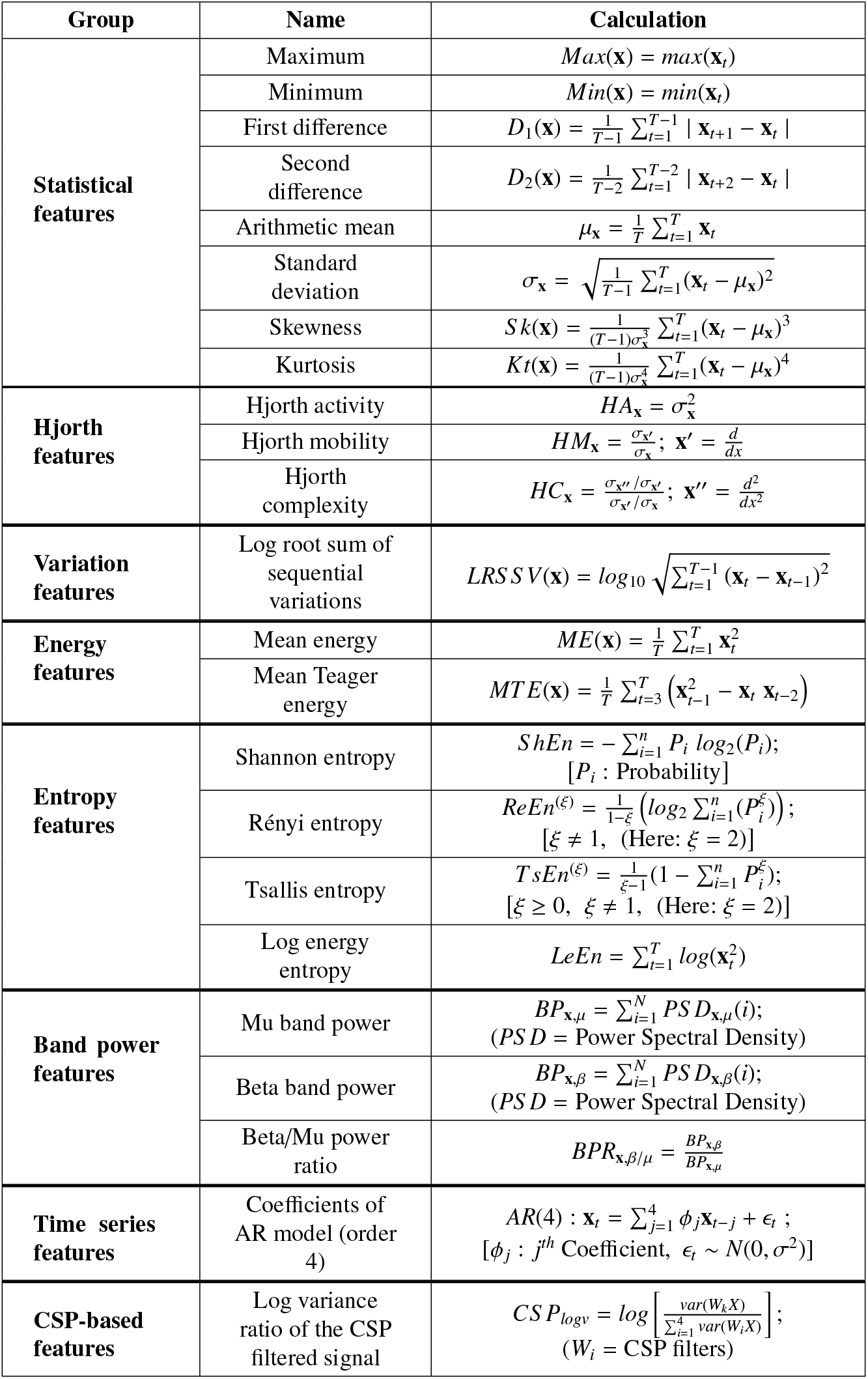
Description of classical metric considered in the bag of features used for MI decoding. A subset of these classical features in combination with GSP features were selected (cf. Section 3.2) through differential evolution (cf. Section 2.5 for MI decoding. (**x** = denotes the timeseries associated to an electrode.)

**Fig. S5:**
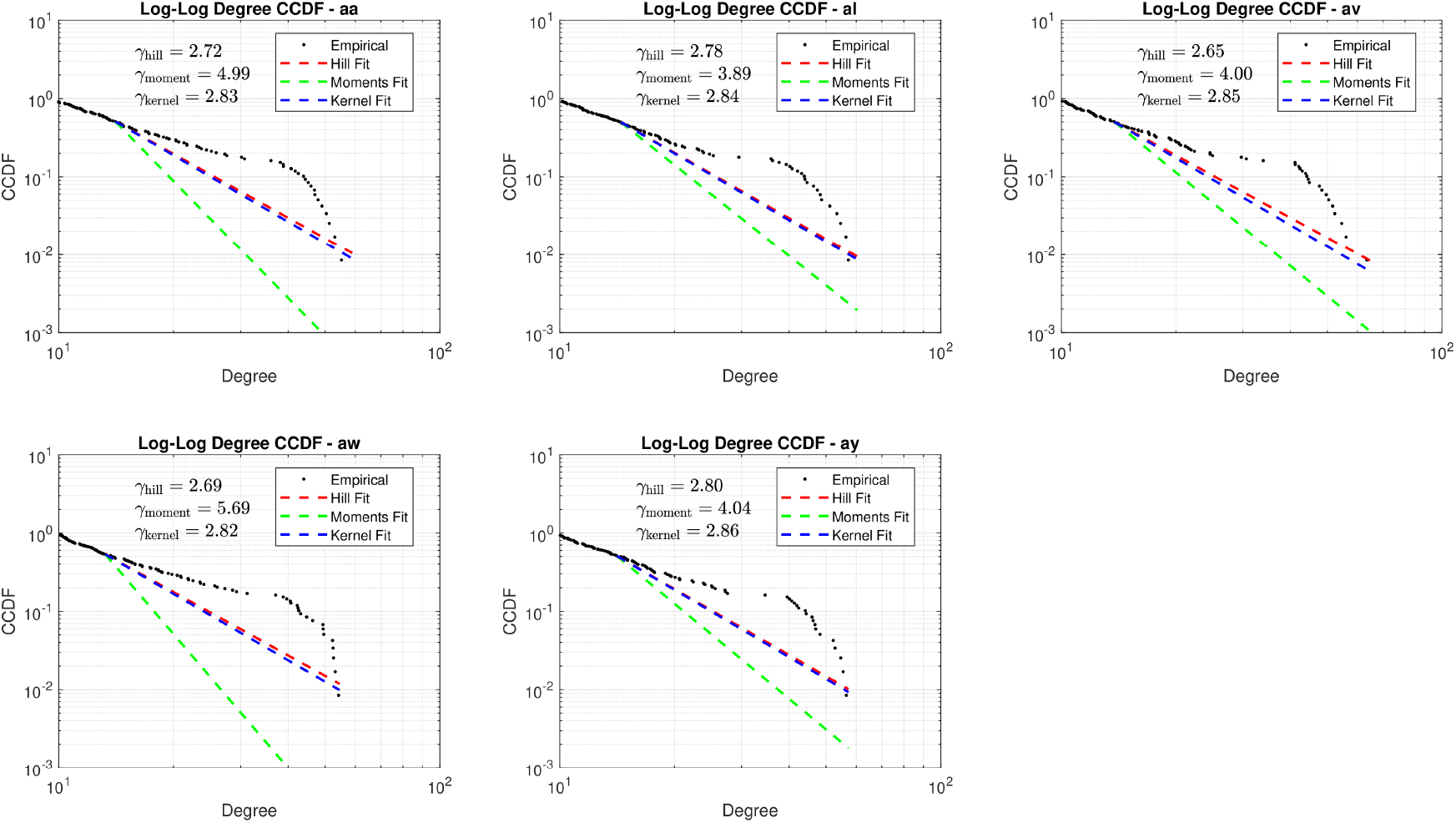
Log-Log CCDFs of degree distributions for five subjects in MI dataset, overlaid with tail exponent estimates from three EVT-based methods: Hill (red), Moment (green), and Kernel (blue). Estimated *γ* values reflect heavy-tailed, scale-free–like structure in brain graphs.

**Table S4:** Paired t-test results (*t*-value and *p*-value) with Bonferroni correction for different modes of brain graph across all subjects. (Significance level: *α*_adj_ = 0.00167.)

| Comparison | $\vert t\vert $-value | $p$-value (raw) | $p$-value (adjusted) |
| --- | --- | --- | --- |
| **Subject aa** |  |  |  |
| SWSF vs PLV | 7.18 | 5.19e-05 | 3.12e-04 |
| SWSF vs SW | 9.45 | 5.72e-06 | 3.43e-05 |
| SWSF vs SF | 6.02 | 1.98e-04 | 1.19e-03 |
| **Subject al** |  |  |  |
| SWSF vs PLV | 8.12 | 1.96e-05 | 1.18e-04 |
| SWSF vs SW | 9.77 | 4.34e-06 | 2.60e-05 |
| SWSF vs SF | 10.85 | 1.81e-06 | 1.08e-05 |
| **Subject av** |  |  |  |
| SWSF vs PLV | 12.31 | 6.19e-07 | 3.72e-06 |
| SWSF vs SW | 10.91 | 1.73e-06 | 1.04e-05 |
| SWSF vs SF | 7.01 | 6.26e-05 | 3.75e-04 |
| **Subject aw** |  |  |  |
| SWSF vs PLV | 11.62 | 1.01e-06 | 6.07e-06 |
| SWSF vs SW | 9.02 | 8.38e-06 | 5.03e-05 |
| SWSF vs SF | 6.45 | 1.18e-04 | 7.09e-04 |
| **Subject ay** |  |  |  |
| SWSF vs PLV | 16.81 | 4.18e-08 | 2.51e-07 |
| SWSF vs SW | 10.41 | 2.56e-06 | 1.54e-05 |
| SWSF vs SF | 5.50 | 3.80e-04 | 2.28e-03 |

**Fig. S6:**
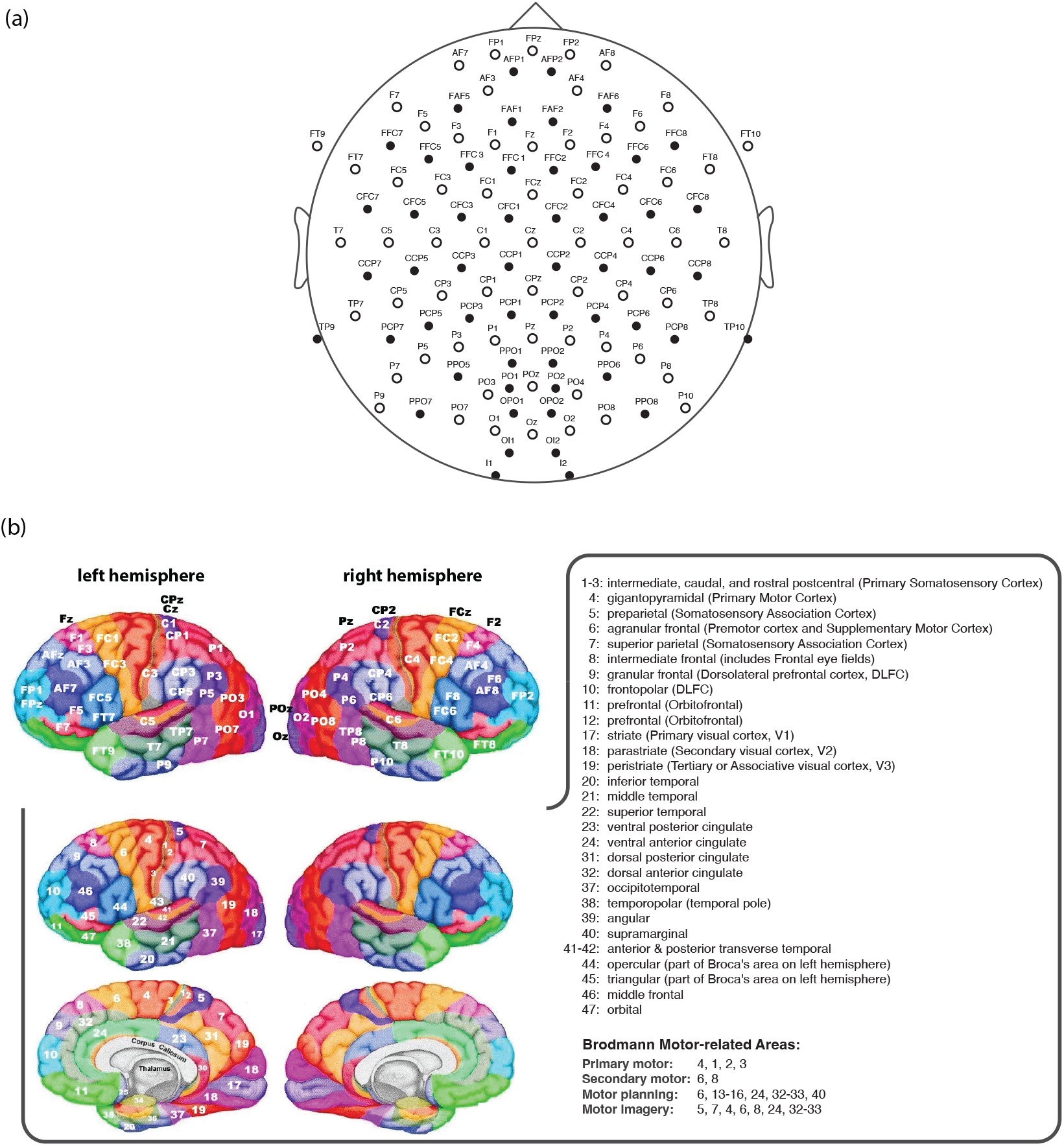
(a) Arrangement of 118 EEG electrodes. Labels, corresponding to cortical locations: frontal (F), temporal (T), central (C), parietal (P), occipital (O), Inion (I). The letter ‘z’ indicates the midline and letter combination refer to intermediate cortical regions. Odd and even numbers indicate the left and right hemisphere, respectively. (b) The position of a subset (64) of the electrodes (marked as white in (a)) are shown on a rendering of the cerebral cortex (top left) relative to Brodmann area centers (bottom left). The surface maps were adopted from http://www.brainm.com/software/pubs/dg/BA_10-20_ROI_Talairach/nearesteeg.html.

**Fig. S7:**
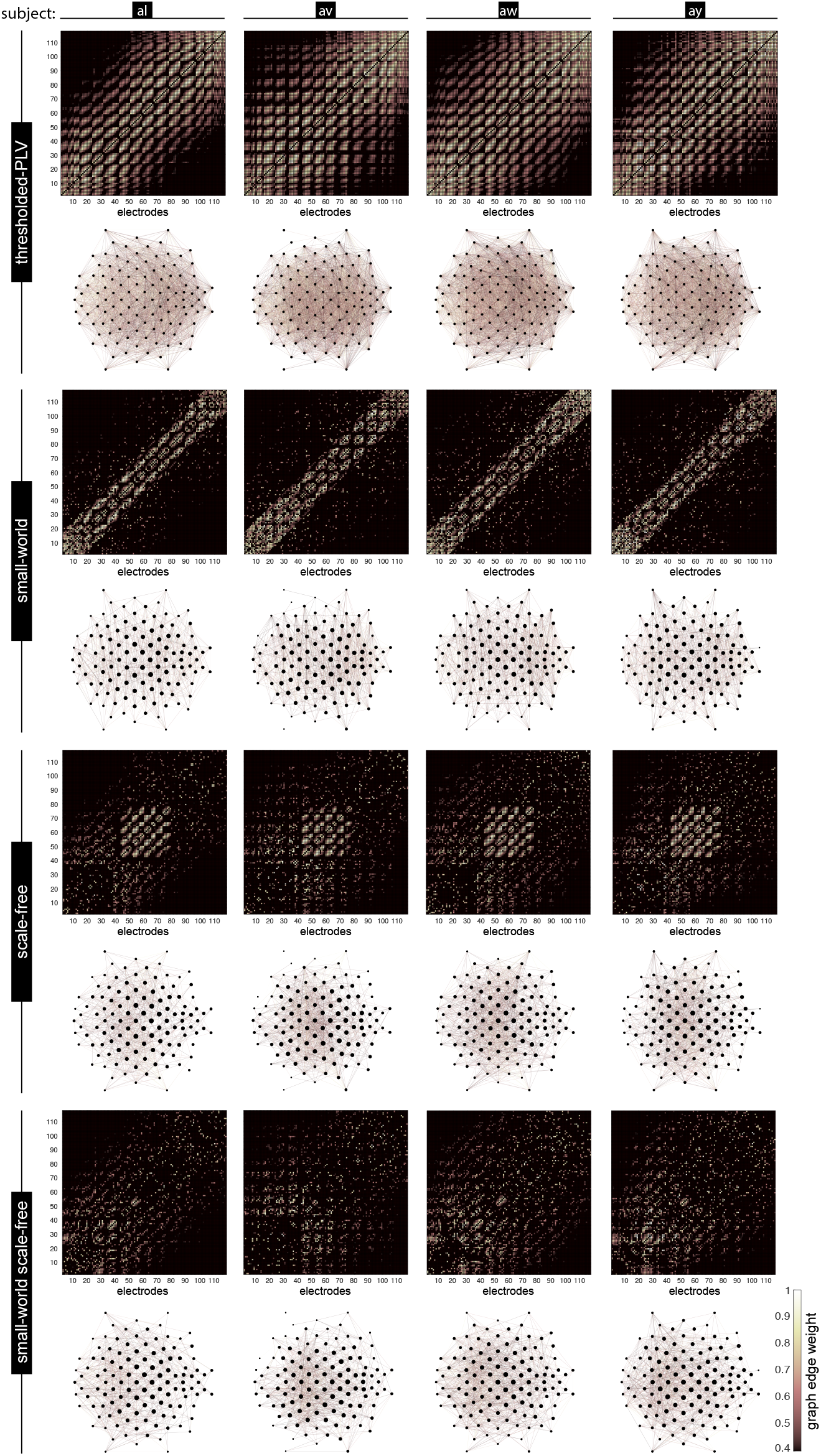
Brain graphs derived from PLV (thresholded), SW, SF, and SWSF models, along with their corresponding weighted adjacency matrices shown for subjects al, av, aw, and ay.

**Table S5:** Paired t-test results (*t*-value and *p*-value) with Bonferroni correction for different Kron-reduced brain graph models across all subjects. (Significance level: *α*_adj_ = 0.00167.)

| Comparison | $\vert t\vert $-value | $p$-value (raw) | $p$-value (adjusted) |
| --- | --- | --- | --- |
| **Subject aa** |  |  |  |
| SWSF<sub>Kron</sub> vs PLV<sub>Kron</sub> | 7.05 | 5.99e−05 | 3.59e−04 |
| SWSF<sub>Kron</sub> vs SW<sub>Kron</sub> | 9.30 | 6.52e−06 | 3.91e−05 |
| SWSF<sub>Kron</sub> vs SF<sub>Kron</sub> | 5.90 | 2.29e−04 | 1.37e−03 |
| **Subject al** |  |  |  |
| SWSF<sub>Kron</sub> vs PLV<sub>Kron</sub> | 8.45 | 1.43e−05 | 8.56e−05 |
| SWSF<sub>Kron</sub> vs SW<sub>Kron</sub> | 9.95 | 3.73e−06 | 2.24e−05 |
| SWSF<sub>Kron</sub> vs SF<sub>Kron</sub> | 11.21 | 1.37e−06 | 8.23e−06 |
| **Subject av** |  |  |  |
| SWSF<sub>Kron</sub> vs PLV<sub>Kron</sub> | 12.02 | 7.59e−07 | 4.55e−06 |
| SWSF<sub>Kron</sub> vs SW<sub>Kron</sub> | 10.61 | 2.18e−06 | 1.31e−05 |
| SWSF<sub>Kron</sub> vs SF<sub>Kron</sub> | 6.85 | 7.47e−05 | 4.48e−04 |
| **Subject aw** |  |  |  |
| SWSF<sub>Kron</sub> vs PLV<sub>Kron</sub> | 11.12 | 1.47e−06 | 8.81e−06 |
| SWSF<sub>Kron</sub> vs SW<sub>Kron</sub> | 8.80 | 1.03e−05 | 6.15e−05 |
| SWSF<sub>Kron</sub> vs SF<sub>Kron</sub> | 6.30 | 1.41e−04 | 8.46e−04 |
| **Subject ay** |  |  |  |
| SWSF<sub>Kron</sub> vs PLV<sub>Kron</sub> | 15.41 | 8.92e−08 | 5.35e−07 |
| SWSF<sub>Kron</sub> vs SW<sub>Kron</sub> | 10.02 | 3.52e−06 | 2.11e−05 |
| SWSF<sub>Kron</sub> vs SF<sub>Kron</sub> | 5.30 | 4.90e−04 | 2.96e−03 |

**Table S6:** MI decoding accuracy (mean ± std,%) of the proposed method for six different models of brain graph (*N* = 118, *M* = 1180) using a cross-subject paradigm. For CLA, the top 10 classical features were used whereas for PLV, SW, SF, and SWSF the top 10 GSP-based and classical features were used. Moreover, the results of EEGNet are shown as a representative lightweight deep learning architecture. The symbol * indicates that the SWSF model is significantly better than all other models, paired t-test, Bonferroni-corrected, *α*_*ad j*_ = 0.00167, p < 0.00167).

| Subject | CLA | PLV | SW | SF | SWSF | EEGNet |
| --- | --- | --- | --- | --- | --- | --- |
| aa | 68.93±3.78 | 70.36±2.41 | 74.29±3.69 | 72.14±4.70 | **78.21±3.55\*** | 61.43±3.28 |
| al | 81.79±2.03 | 82.50±4.60 | 87.14±1.84 | 84.64±3.39 | **90.71±3.01\*** | 88.57±2.82 |
| av | 60.71±3.76 | 62.14±5.11 | 67.86±3.76 | 63.93±3.55 | **72.14±4.05\*** | 54.29±3.69 |
| aw | 71.79±3.55 | 72.86±3.84 | 78.57±2.38 | 75.71±2.82 | **86.07±2.64\*** | 65.36±4.78 |
| ay | 70.36±2.94 | 71.43±5.05 | 77.86±2.82 | 75.00±4.12 | **85.36±2.64\*** | 62.50±3.86 |
| mean±std | 70.71±7.53 | 71.79±7.26 | 77.14±7.02 | 74.29±7.44 | **82.50±7.31\*** | 66.43±13.03 |

**Table S7:** MI decoding accuracy (mean ± std,%) of the proposed method for four different models of brain graph with Kron reduction using a cross-subject paradigm. The symbol * indicates that the SWSF_Kron_ model is significantly better than all other models (paired t-test, Bonferroni-corrected, *α*_*ad j*_ = 0.00167, p < 0.00167).

| Subject | PLV<sub>Kron</sub> | SW<sub>Kron</sub> | SF<sub>Kron</sub> | SWSF<sub>Kron</sub> |
| --- | --- | --- | --- | --- |
| aa | 72.14±3.28 | 75.71±4.05 | 74.29±3.28 | **81.79±3.13\*** |
| al | 83.93±1.88 | 88.93±3.13 | 86.79±2.41 | **92.50±2.64\*** |
| av | 64.29±4.12 | 71.07±3.55 | 67.14±3.69 | **75.36±3.93\*** |
| aw | 75.00±2.92 | 81.43±2.26 | 78.57±2.38 | **88.57±3.28\*** |
| ay | 74.29±3.28 | 81.07±2.94 | 77.14±3.45 | **88.93±3.55\*** |
| mean±std | 73.93±7.03 | 79.64±6.71 | 76.79±7.12 | **85.36±6.83\*** |

## Footnotes

1 Different methods can be used to estimate the analytic signal, here, we used the fast Fourier transform-based implementation [54] from MATLAB.

